# DiffDomain-Spectrum identifies structurally reorganized TADs from sparse aggregated single-cell Hi-C contact maps

**DOI:** 10.64898/2026.09.15.751728

**Authors:** Jiadi Zhu, Haishan Zhang, Yanyi Du, Xiao Zhang, Yan Zhou, Dechao Tian

## Abstract

Structurally reorganized topologically associating domains (TADs) capture condition- or cell-type-specific remodeling of chromatin contacts and are important for understanding genome organization in health and disease. Emerging single-cell Hi-C (scHi-C) technologies enable such comparisons across heterogeneous cell populations, but aggregated scHi-C contact maps remain sparse at biologically meaningful 25 kb resolution, limiting reliable TAD reorganization detection. Here we present DiffDomain-Spectrum, a spectral statistical framework for identifying reorganized TADs between conditions or cell types from aggregated raw scHi-C contact maps. It tests normalized TAD-level difference matrices without separately normalizing sparse maps or enhancing individual scHi-C contact maps. Comparison with a semicircle-law null integrates evidence across the full eigenvalue spectrum. Across multiple scHi-C platforms, DiffDomain-Spectrum balances false positive control and detection sensitivity relative to alternative bulk callers, and detects a substantially higher proportion of reference TADs as reorganized than the boundary-focused single-cell method scHiCluster. Detected TADs show coherent aggregate contact patterns and CTCF binding changes and are enriched for differentially expressed genes, supporting biological relevance. Together, these results establish DiffDomain-Spectrum as a statistically principled framework for comparative domain-level analysis of sparse aggregated scHi-C contact maps without single-cell map enhancement.

## Introduction

High-throughput chromosome conformation capture (Hi-C) revealed that mammalian genomes are organized into topologically associating domains (TADs), contiguous genomic regions characterized by preferential chromatin interactions within domains and insulation across domain boundaries [1–5]. TADs are reorganized across biological conditions and disease contexts, providing a structural basis for linking condition-specific 3D genome changes to regulatory function [6–11]. Single-cell Hi-C (scHi-C) measures chromatin contacts in individual cells, enabling cell-type-resolved analysis of 3D genome organization across complex tissues and biological conditions [12–16]. scHi-C therefore enables comparative analysis of 3D genome organization across cell types and states within heterogeneous samples, rather than only across bulk maps averaged over such samples [16–18]. Recent scHi-C studies have linked cell-type-specific chromatin architecture to cellular identity, developmental progression and disease-related states, revealing differences in TAD-like boundaries, domain organization and chromosome conformation [16, 19–22]. Together, these advances establish scHi-C data as a foundation for detecting condition- and cell-type-specific TAD reorganization, thereby connecting 3D genome structure with regulatory function in complex tissues.

Realizing this potential, however, requires overcoming the extreme sparsity of scHi-C contact maps, particularly at the high resolutions required for comparative domain analysis [23, 24]. One strategy is to enhance individual scHi-C contact maps before comparison. The boundary-focused single-cell method scHiCluster, for example, enhances each contact map before testing differences in TAD boundary occurrence between cell populations [25]. Although this approach yields interpretable boundary-level comparisons, boundary-focused testing can omit reorganized TADs without boundary changes, which accounted for a median of 43% of reorganized TADs across bulk Hi-C comparisons [4, 26]. More broadly, enhancement and representation-learning workflows become increasingly computationally demanding as genomic resolution and cell number increase [22, 25, 27–29]. In a recent benchmark, the scHiCluster workflow exceeded 256 GB of memory at 100 kb resolution for 1,800 cells and at 200 kb or finer for 18,000 cells, at resolutions still substantially coarser than the 25 kb used here [29]. For cell-type-level comparison, aggregating raw contacts across cells of the same type or condition offers a scalable route that improves coverage without per-cell enhancement and enables existing bulk Hi-C methods to be applied [16, 18–21, 30–33]. Compared with conventional bulk Hi-C maps, however, aggregated scHi-C contact maps remain substantially lower in coverage and more sparse, with a median of 29.99% (IQR: 24.91%–38.42%) of bin-pair entries within 2 Mb containing zero observed contacts at 25 kb resolution across aggregated maps. Their coverage can also differ substantially across cell types or conditions owing to unequal cell numbers, with up to 14.22-fold differences in cell numbers across aggregated maps [20]. These sparsity and coverage properties, which distinguish aggregated scHi-C contact maps from conventional bulk Hi-C maps, define a data regime for which existing bulk Hi-C callers were not specifically developed. Together, enhancement-based single-cell methods and bulk methods applied to aggregated maps have expanded comparative scHi-C analysis, but they leave unresolved the need for computational methods tailored to domain-level comparison of sparse aggregated scHi-C contact maps. Here we present DiffDomain-Spectrum, a spectral statistical framework for identifying structurally reorganized TADs between conditions or cell types from high-resolution, typically 25 kb, aggregated raw scHi-C contact maps. DiffDomain-Spectrum begins with a reference TAD set, typically called from the condition-1 aggregated contact map using the insulation-score method, and extracts matched TAD-level contact matrices from the two aggregated maps. It constructs normalized TAD-level difference matrices through log-scale comparison, distance-stratified *z*-score standardization and sparsity-aware stabilization of missing and extreme entries, without separately normalizing sparse maps or enhancing individual scHi-C contact maps. The full eigenvalue spectrum of each normalized difference matrix is compared with a semicircle-law null, integrating evidence across the spectrum, and TAD-specific significance is calibrated by Monte Carlo sampling from the semicircle distribution rather than by resampling cells or reconstructing contact maps. Although motivated by the spectral testing principle underlying our earlier bulk Hi-C method DiffDomain [26], DiffDomain-Spectrum develops a distinct full-spectrum testing framework for the low-coverage and sparse regime of aggregated scHi-C contact maps. Across scHi-C datasets generated using multiple platforms, DiffDomain-Spectrum balances false positive control and detection sensitivity relative to alternative bulk callers and detects a substantially higher proportion of reference TADs as reorganized than the boundary-focused single-cell method scHiCluster. Detected reorganized TADs show subtype-consistent aggregate contact patterns and CTCF binding changes and are enriched for differentially expressed genes, supporting their biological relevance. Together, these results establish DiffDomain-Spectrum as a statistically principled framework for comparative domain-level analysis of sparse aggregated scHi-C contact maps without single-cell map enhancement.

## Results

### Overview of DiffDomain-Spectrum

DiffDomain-Spectrum identifies structurally reorganized TADs between two biological conditions from sparse aggregated scHi-C contact maps (Fig. 1). For each condition, raw scHi-C contacts from cells of the same cell type or condition are aggregated into contact maps, improving effective contact coverage without enhancing individual scHi-C contact maps (Fig. 1a). Given a reference TAD list, typically defined from the aggregated contact map of condition 1 using the insulation score method [34], DiffDomain-Spectrum extracts the corresponding *N ×N* contact submatrices from the two conditions for each TAD, denoted *M*_1_ and *M*_2_ (Fig. 1b). It then constructs a log-scale TAD-level difference matrix, stabilizes sparse and extreme entries, and applies distance-stratified *z*-score normalization to reduce systematic effects associated with genomic distance, sequencing depth and unequal cell numbers between conditions. The resulting normalized difference matrix, *M*_norm_, summarizes TAD-level contact-pattern differences after reducing global and distance-dependent effects (Fig. 1c).

**Figure 1:**
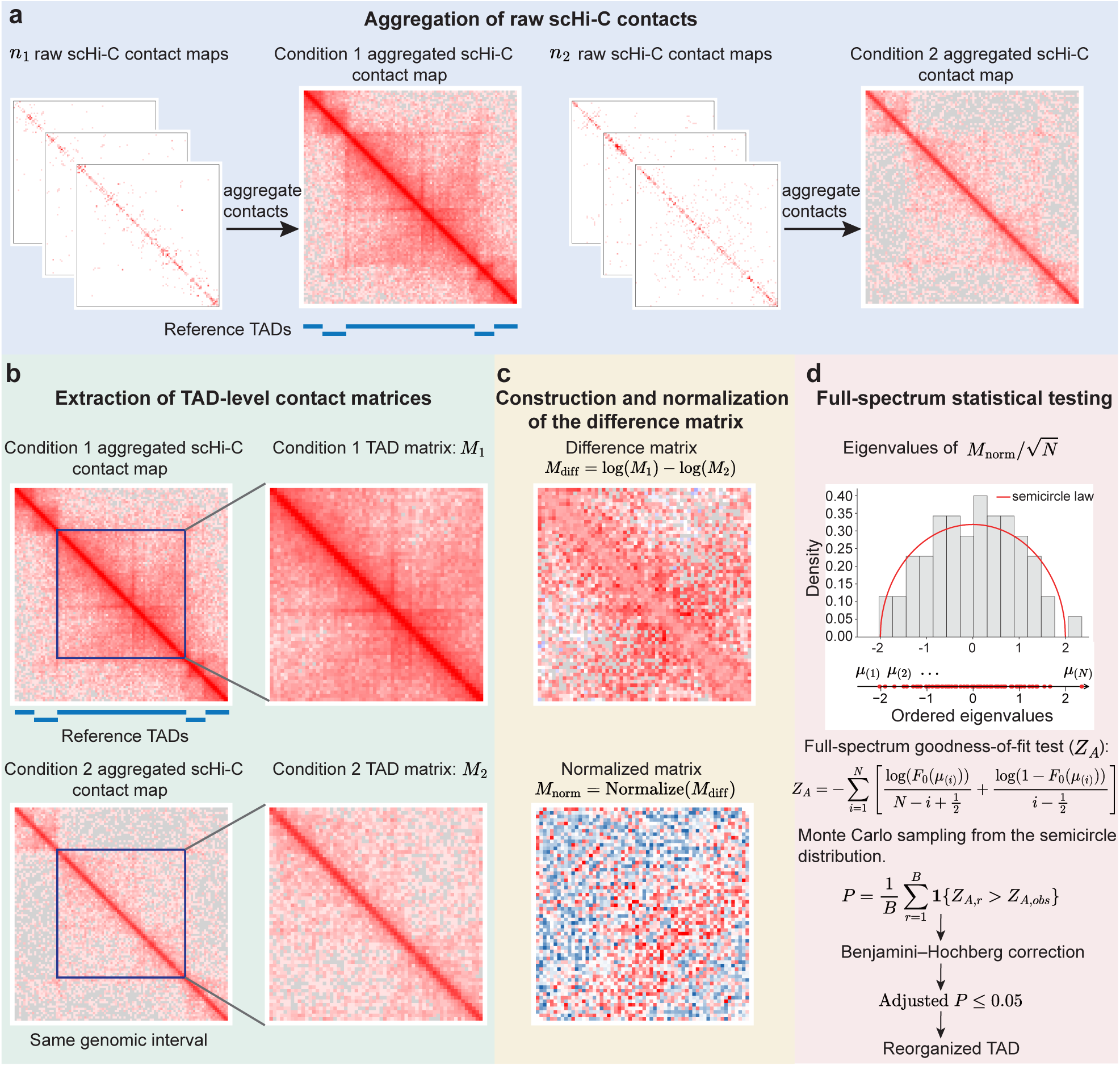
Workflow of DiffDomain-Spectrum for identifying structurally reorganized TADs from sparse aggregated scHi-C contact maps. **a**, Raw scHi-C contact maps from *n*_1_ and *n*_2_ cells are aggregated separately within conditions 1 and 2 to generate aggregated scHi-C contact maps, without enhancing individual scHi-C contact maps. Reference TADs are defined by default from the condition-1 aggregated contact map using the insulation score. **b**, The same two aggregated contact maps from panel a are used for TAD-level comparison. For each condition-1 reference TAD, the corresponding genomic interval is extracted from both aggregated maps to obtain the *N ×N* contact matrices *M*_1_ and *M*_2_. **c**, A log-scale TAD-level difference matrix is constructed as *M*_diff_ = log(*M*_1_) − log(*M*_2_) and normalized to obtain *M*_norm_ = Normalize(*M*_diff_). Normalization includes missing and infinite-value stabilization, distance-specific standardization, all-missing-offset replacement, and clipping. **d**, The full eigenvalue spectrum of 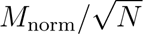 is compared with the semicircle-law null distribution using the *Z_A_*full-spectrum goodness-of-fit test. Statistical significance is estimated by TAD-specific Monte Carlo sampling from the semicircle distribution conditional on the observed spectral range, followed by Benjamini–Hochberg correction across tested TADs. All contact maps, TAD-level matrices, difference and normalized matrices, and the eigenvalue distribution shown in panels a–d are derived from GM12878–K562 comparison over chr11:20150000–22575000 using LiMCA dataset [51]. The displayed TAD (chr11:20650000– 22125000) is detected as reorganized.

DiffDomain-Spectrum formulates reorganized TAD detection as a spectral goodness-of-fit test. Under the null hypothesis of no structural reorganization, the spectrum of 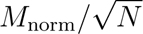 is modeled using an approximate generalized-Wigner spectral null [35, 36], for which the semicircle law provides the limiting reference distribution [37]. DiffDomain-Spectrum therefore compares the empirical eigenvalue distribution of 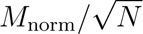, denoted *F_N_*, with the theoretical cumulative distribution function of the semicircle law, denoted *F*_0_. Under the spectral null of no structural reorganization, *F_N_*is expected to be compatible with *F*_0_, whereas systematic departure from *F*_0_ provides evidence of structural reorganization. In contrast to DiffDomain, which focuses on the largest eigenvalue [26], DiffDomain-Spectrum uses the full eigenvalue spectrum to capture coordinated contact-pattern changes that are distributed across sparse aggregated scHi-C contact maps (Fig. 1d). The deviation between *F_N_* (*x*) and *F*_0_(*x*) is quantified using a full-spectrum test based on the likelihood-ratio goodness-of-fit statistic *Z_A_* [38]; statistical significance is estimated by Monte Carlo sampling from the semicircle distribution conditional on the observed spectral range, avoiding repeated cell subsampling or contact-map reconstruction. For each TAD, DiffDomain-Spectrum computes a *P*-value and applies Benjamini–Hochberg correction across tested TADs; TADs with adjusted *P* ≤ 0.05 are identified as significantly reorganized and used in subsequent benchmarking and biological analyses.

### DiffDomain-Spectrum balances false positive control and detection sensitivity

We first evaluated whether DiffDomain-Spectrum controls false positives in comparisons for which no systematic TAD reorganization was expected. Empirical false positive rates (FPRs) were estimated from ten control comparisons between datasets representing the same biological condition, treating all detected reorganized TADs as false positives. These comprised three biological-replicate comparisons from mouse cortex Dip-C data [15]; four mouse embryonic stem cell (mESC) dscHi-C controls, including one biological-replicate comparison, two replicate-versus-subsample comparisons and one comparison between biotin and no-biotin preparations [39]; and three pairwise comparisons among GM12878 biological replicates profiled by scNanoHi-C2 [40]. Dip-C and dscHi-C data were analyzed at 25 kb resolution, whereas scNanoHi-C2 data were analyzed at 50 kb resolution. We compared DiffDomain-Spectrum with the alternative bulk callers DiffDomain, DiffGR, TADCompare, DiffTAD-parametric, DiffTAD-permutation, HiCcompare and HiCDC+, with HiCcompare and HiCDC+ adapted here for reorganized TAD detection [26, 41–45]. To ensure comparability, all methods were applied to the same aggregated scHi-C contact maps and evaluated using Benjamini–Hochberg-adjusted *P*-values at a significance threshold of 0.05.

At a representative genomic region encompassing *Sox6* in mESCs (chr7:114,000,000–117,500,000), the two replicate aggregated contact maps showed similar domain-level contact patterns. Consistently, DiffDomain-Spectrum, DiffDomain and HiCcompare did not detect significant TAD reorganization in this region (Fig. 2a). By contrast, several alternative bulk callers identified reorganized TADs, with DiffTAD-permutation and HiCDC+ classifying more than half of the TADs in this region as significant, indicating inflated false positive detection. Genome-wide analysis showed a similar pattern (Fig. 2b). DiffDomain-Spectrum, DiffDomain and HiCcompare maintained empirical FPRs below 5%, whereas DiffTAD-parametric, DiffTAD-permutation and HiCDC+ showed substantially inflated FPRs exceeding 50%; DiffGR and TADCompare also exceeded the nominal threshold. Across all ten control comparisons spanning three scHi-C platforms, DiffDomain-Spectrum and DiffDomain consistently maintained low and stable FPRs, whereas each of the remaining methods exceeded the nominal 5% level in one or more controls and showed greater variation in FPR across control comparisons (Fig. 2c). These results indicate that DiffDomain-Spectrum consistently controls the empirical FPR across sparse aggregated scHi-C contact maps generated using diverse platforms.

**Figure 2:**
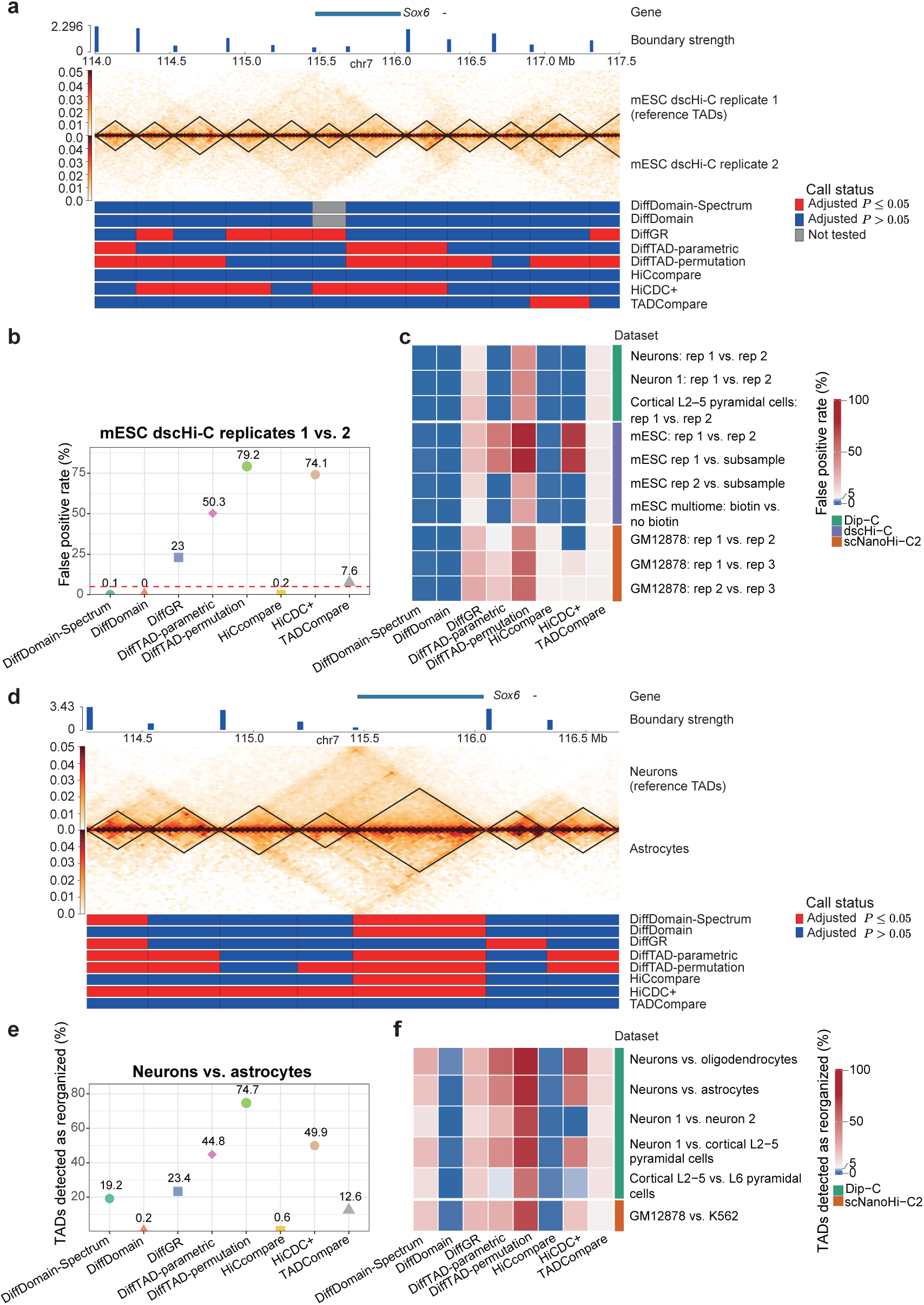
Benchmarking false positive control and detection sensitivity of DiffDomain-Spectrum against alternative bulk callers. **a**, Locus-level reorganized-TAD calls in a representative genomic region encompassing *Sox6* (chr7:114,000,000–117,500,000) for two aggregated mESC dscHi-C replicate contact maps. TAD boundaries defined from replicate 1 are overlaid on both contact maps and constitute the reference TAD set. Rows below the maps show the call status assigned to each reference TAD by the eight methods. Red indicates TADs detected as reorganized at Benjamini–Hochberg-adjusted *P ≤* 0.05, blue indicates TADs not detected as reorganized at adjusted *P >* 0.05, and gray indicates TADs that were not tested because they contained fewer than 10 genomic bins and were excluded by DiffDomain-Spectrum and DiffDomain. **b**, Genome-wide empirical false positive rates (FPRs) obtained by comparing aggregated mESC dscHi-C replicate 1 and replicate 2 contact maps. The red dashed line denotes the nominal 5% FPR level. **c**, Empirical FPRs across ten controlcomparisons for which no systematic TAD reorganization is expected, comprising three mouse cortex Dip-C comparisons, four mESC dscHi-C comparisons and three GM12878 scNanoHi-C2 comparisons. Dip-C and dscHi-C data were analyzed at 25 kb resolution, and scNanoHi-C2 data were analyzed at 50 kb resolution. The color scale is centered at the nominal 5% level. **d**, Locus-level reorganized-TAD calls in a representative genomic region encompassing *Sox6* (chr7:114,250,000–116,650,000) in the neurons-versus-astrocytes comparison. TAD boundaries defined from the aggregated neurons contact map are overlaid on both maps and constitute the reference TAD set. Rows below the maps show the call status assigned to each reference TAD by the eight methods, with red and blue indicating adjusted *P* ≤ 0.05 and adjusted *P >* 0.05, respectively. **e**, Genome-wide proportion of neurons-defined reference TADs detected as reorganized by each method in the neurons-versus-astrocytes comparison. **f**, Proportions of reference TADs detected as reorganized across all six biological comparisons, comprising five mouse cortical cell-type comparisons profiled by Dip-C at 25 kb resolution and the GM12878-versus-K562 comparison profiled by scNanoHi-C2 at 50 kb resolution. The neurons-versus-astrocytes comparison shown separately in **e** is included in this summary.

We next assessed detection sensitivity in comparisons between distinct biological conditions. This analysis included five comparisons between mouse cortical cell types profiled by Dip-C and one comparison between human GM12878 and K562 cell lines profiled by scNanoHi-C2, analyzed at 25 kb and 50 kb resolution, respectively. The same alternative bulk callers were evaluated as in the FPR analysis. We again examined a representative genomic region surrounding *Sox6* in the neurons-versus-astrocytes comparison (chr7:114,250,000–116,650,000). *Sox6* is a transcription factor involved in neuronal differentiation and cell fate specification and exhibits cell-type-specific functions in the brain [46–48]. Six of the eight methods identified the TAD containing *Sox6* as reorganized, whereas DiffGR and TADCompare did not detect significant reorganization (Fig. 2d). The established cell-type-specific functions of *Sox6* support the biological plausibility of the detected reorganization at this locus, although this locus-level observation does not provide an independent measure of method accuracy.

At the genome-wide level, DiffDomain-Spectrum detected an intermediate proportion of reorganized TADs relative to the alternative bulk callers (Fig. 2e). DiffTAD-permutation reported the highest detection proportion, identifying 74.7% of TADs as reorganized, while DiffTAD-parametric and HiCDC+ each identified nearly half of all TADs. These high detection proportions cannot be interpreted as evidence of greater detection sensitivity because the same methods exhibited markedly inflated FPRs in control comparisons. By contrast, DiffDomain and HiCcompare detected fewer than 5% of TADs, indicating conservative detection that may reduce sensitivity. Across additional datasets generated using different scHi-C protocols, DiffDomain-Spectrum generally maintained intermediate detection proportions, whereas the DiffTAD variants and HiCDC+ detected larger fractions of TADs and DiffDomain and HiCcompare remained conservative (Fig. 2f). Thus, DiffDomain-Spectrum maintained moderate detection sensitivity without the marked FPR inflation observed for methods with higher detection proportions.

To assess whether detected reorganized TADs exhibited subtype-consistent domain-level contact changes, we performed aggregated domain analysis (ADA), which summarizes average contact patterns across detected TADs. Reorganized TADs detected by each method were assigned to six structural subtypes following the DiffDomain framework: *strength-change*, *loss*, *zoom*, *merge*, *split* and *complex* (Methods). The *strength-change* subtype was further separated into *strength-change up* and *strength-change down*, with detailed subtype definitions provided in Methods. Applying the same subtype-classification procedure to all methods enabled direct comparison of the aggregate contact patterns associated with each assigned subtype. To ensure that each ADA profile was supported by a sufficient number of reorganized TADs, we restricted the analysis to methods that detected more than 5% of TADs as reorganized and excluded the *loss* subtype because the number of TADs assigned to this subtype was insufficient for reliable aggregation. For the representative *strength-change up* subtype, DiffDomain-Spectrum produced aggregate contact enrichment confined within the expected domain region, consistent with the subtype definition (Supplementary Fig. 1a,b). By contrast, DiffGR, DiffTAD-parametric, DiffTAD-permutation and TADCompare produced enrichment patterns extending beyond the expected TAD boundaries, suggesting that some detected regions may reflect boundary shifts or spatially diffuse contact changes rather than localized changes in intra-domain interaction strength. Similar concordance between DiffDomain-Spectrum ADA profiles and the corresponding subtype definitions was observed for the other evaluated reorganization subtypes (Supplementary Fig. 1c–g). Overall, the ADA profiles indicate that reorganized TADs detected by DiffDomain-Spectrum exhibit, on average, greater concordance with the expected subtype-specific contact patterns than those detected by the evaluated alternative bulk callers.

Taken together, DiffDomain-Spectrum was the only method among those evaluated that consistently combined low empirical FPRs across the ten controls with moderate detection sensitivity across comparisons between distinct biological conditions or cell types. Its subtype-consistent ADA profiles further support the reliable identification of structurally coherent TAD reorganization from sparse aggregated scHi-C contact maps.

### DiffDomain-Spectrum detects substantially more reorganized TADs than the boundary-focused single-cell method scHiCluster

We compared DiffDomain-Spectrum with the boundary-focused single-cell method scHiCluster [25, 49]. scHiCluster applies TopDom separately to each enhanced individual scHi-C contact map to identify TAD boundaries, quantifies boundary occurrence across cells and tests differences between cell populations to identify differential boundaries. DiffDomain-Spectrum instead evaluates contact-pattern differences within reference TADs using aggregated raw scHi-C contact maps, without enhancing individual scHi-C contact maps. We analyzed all 30 directed pairwise comparisons among six cell types from developing mouse embryos profiled by HiRES [20]. For each comparison, TopDom-defined TADs from the condition-1 aggregated scHi-C contact map served as the common TAD-level evaluation set. Because scHiCluster differential boundaries were derived from TopDom calls on enhanced individual maps, whereas the reference TADs were defined from aggregated maps, the differential-boundary coordinates could differ from the boundaries of the reference TADs. A reference TAD was therefore considered reorganized by scHiCluster when at least one differential boundary fell within its genomic interval. This design retained the native scHiCluster workflow while enabling direct comparison on the same reference TAD set.

Across the 30 comparisons, scHiCluster detected a mean of 100 differential boundaries per comparison, with a median of 15 and a maximum of 405. These differential-boundary counts are comparable to those from an independent published application of scHiCluster [50]. After the differential boundaries were mapped to the common reference TAD sets, DiffDomain-Spectrum detected a higher proportion of reference TADs as reorganized in all 30 comparisons. The median detection proportions were 24.77% for DiffDomain-Spectrum and 0.32% for scHiCluster (Fig. 3a,b). scHiCluster tests differential boundary occurrence across individual cells, whereas DiffDomain-Spectrum tests changes across the full contact pattern within each reference TAD in aggregated scHi-C contact maps. The two methods therefore evaluate related but distinct aspects of TAD reorganization.

**Figure 3:**
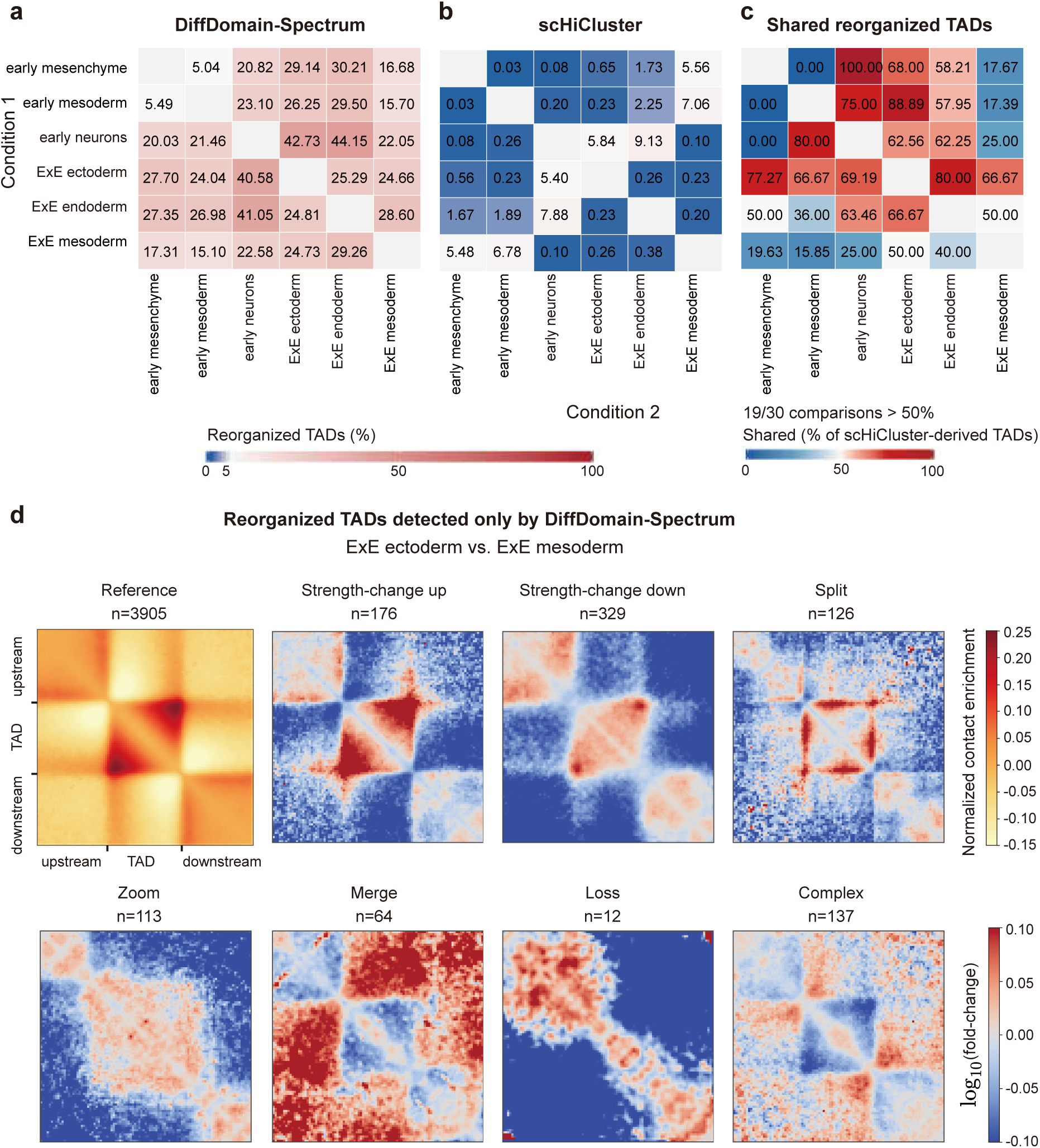
Comparison of DiffDomain-Spectrum and scHiCluster for detecting reorganized TADs across cell types in developing mouse embryos. **a**, Proportion of condition-1 reference TADs detected as reorganized by DiffDomain-Spectrum across 30 directed pairwise comparisons among six cell types. Rows denote condition 1 and columns denote condition 2. DiffDomain-Spectrum was applied to aggregated raw scHi-C contact maps constructed from raw contacts pooled across cells. **b**, Proportion of condition-1 reference TADs considered reorganized based on scHiCluster differential-boundary calls across the same comparisons. **c**, Proportion of reorganized TADs derived from scHiCluster differential-boundary calls that were also detected as reorganized by DiffDomain-Spectrum. **d**, Subtype-level aggregated domain analysis (ADA) of reorganized TADs detected only by DiffDomain-Spectrum in the ExE ectoderm-versus-ExE mesoderm comparison. The leftmost map shows the reference ADA profile of ExE ectoderm-defined TADs. The remaining maps show log_10_(*M*_2_*/M*_1_) for the seven structural subtypes, where *M*_1_ is the reference ADA matrix and *M*_2_ is the subtype-specific ADA matrix generated from the ExE mesoderm aggregated scHi-C contact map. The number of reorganized TADs contributing to each subtype is shown beneath the label. Corresponding condition-2 ADA matrices and fold-change matrices for four representative comparisons are shown in Supplementary Fig. 2.

Despite the large difference in call-set size, DiffDomain-Spectrum captured a substantial fraction of the reorganized TADs derived from scHiCluster differential-boundary calls. In 19 of the 30 comparisons, more than 50% of these reorganized TADs were also identified as reorganized by DiffDomain-Spectrum (Fig. 3c). This result demonstrates substantial directional overlap between the DiffDomain-Spectrum calls and the TAD-level calls derived from scHiCluster differential boundaries.

We next examined the structural composition and aggregate contact patterns of reorganized TADs detected only by DiffDomain-Spectrum. In the ExE ectoderm-versus-ExE mesoderm comparison, 957 reorganized TADs were detected by DiffDomain-Spectrum but not by scHiCluster. These comprised 176 *strength-change up*, 329 *strength-change down*, 126 *split*, 113 *zoom*, 64 *merge*, 12 *loss* and 137 *complex* reorganized TADs (Fig. 3d). The 505 *strength-change* TADs retained their boundaries, whereas the remaining 452 reorganized TADs belonged to the five subtypes involving boundary changes.

Subtype-level aggregated domain analysis averaged contact patterns across reorganized TADs within each subtype. The *strength-change up* and *strength-change down* groups showed increased and decreased within-domain contacts, respectively, while boundary positions were retained. The *split* group showed broadly weaker within-domain contacts, with pronounced reductions near both domain boundaries. The *zoom* group showed redistributed contacts within reference TADs and adjacent regions. The *merge* group showed increased contacts between reference TADs and their flanking regions. The *loss* group showed reduced contrast between within-domain and flanking contacts, consistent with weakened domain insulation. The *complex* group showed spatially distributed changes within reference TADs and across their flanking regions. Across three additional developmental comparisons, all seven subtypes were represented and showed broadly subtype-consistent aggregate patterns. Comparable patterns were obtained when all reorganized TADs detected by DiffDomain-Spectrum were analyzed, including those also classified as reorganized based on scHiCluster differential-boundary calls (Supplementary Fig. 2). Thus, the reorganized TADs detected only by DiffDomain-Spectrum included subtypes with preserved boundaries and subtypes involving boundary changes and showed subtype-consistent aggregate contact patterns.

Together, DiffDomain-Spectrum captured a substantial fraction of the reorganized TADs derived from scHiCluster differential-boundary calls while identifying a substantially larger set with subtype-consistent aggregate contact patterns. DiffDomain-Spectrum therefore provides a complementary domain-level framework to boundary-focused single-cell analysis for identifying reorganized TADs from aggregated raw scHi-C contact maps without enhancing individual scHi-C contact maps.

### DiffDomain-Spectrum is robust across cell numbers and genomic resolutions

We first evaluated how cell number and the resulting data sparsity affected the detection proportion and reproducibility of DiffDomain-Spectrum. Using mouse cortex Dip-C data with a median of approximately 400,000 contacts per cell, we repeatedly subsampled increasing numbers of cells from each cell type and pooled their raw contacts to construct aggregated scHi-C contact maps at 25 kb resolution [15]. Sparsity was defined as the proportion of zero-valued entries within the contact matrices of the tested TADs. Reproducibility was quantified using the Jaccard index between each subsample-derived call set and the corresponding call set obtained using all available cells. The sampling design included balanced settings, in which both conditions were subsampled to the same cell number, and unbalanced settings, in which the cell number in one condition was increased while the other was fixed at its maximum available cell number.

Increasing cell numbers in both conditions increased both the proportion of TADs detected as reorganized and the reproducibility of the resulting call sets (Fig. 4a–c). In the neurons-versus-oligodendrocytes comparison, the mean Jaccard index reached 0.33 with 150 cells per condition. With 250 cells per condition, DiffDomain-Spectrum detected a mean of 26.14% of neurons-defined TADs as reorganized and achieved a mean Jaccard index of 0.52 relative to the call set obtained using all available cells (Fig. 4a). Comparable increases in detection proportion and reproducibility were observed in the neurons-versus-astrocytes and neuron 1-versus-cortical L2–5 pyramidal cells comparisons (Fig. 4b,c).

**Figure 4:**
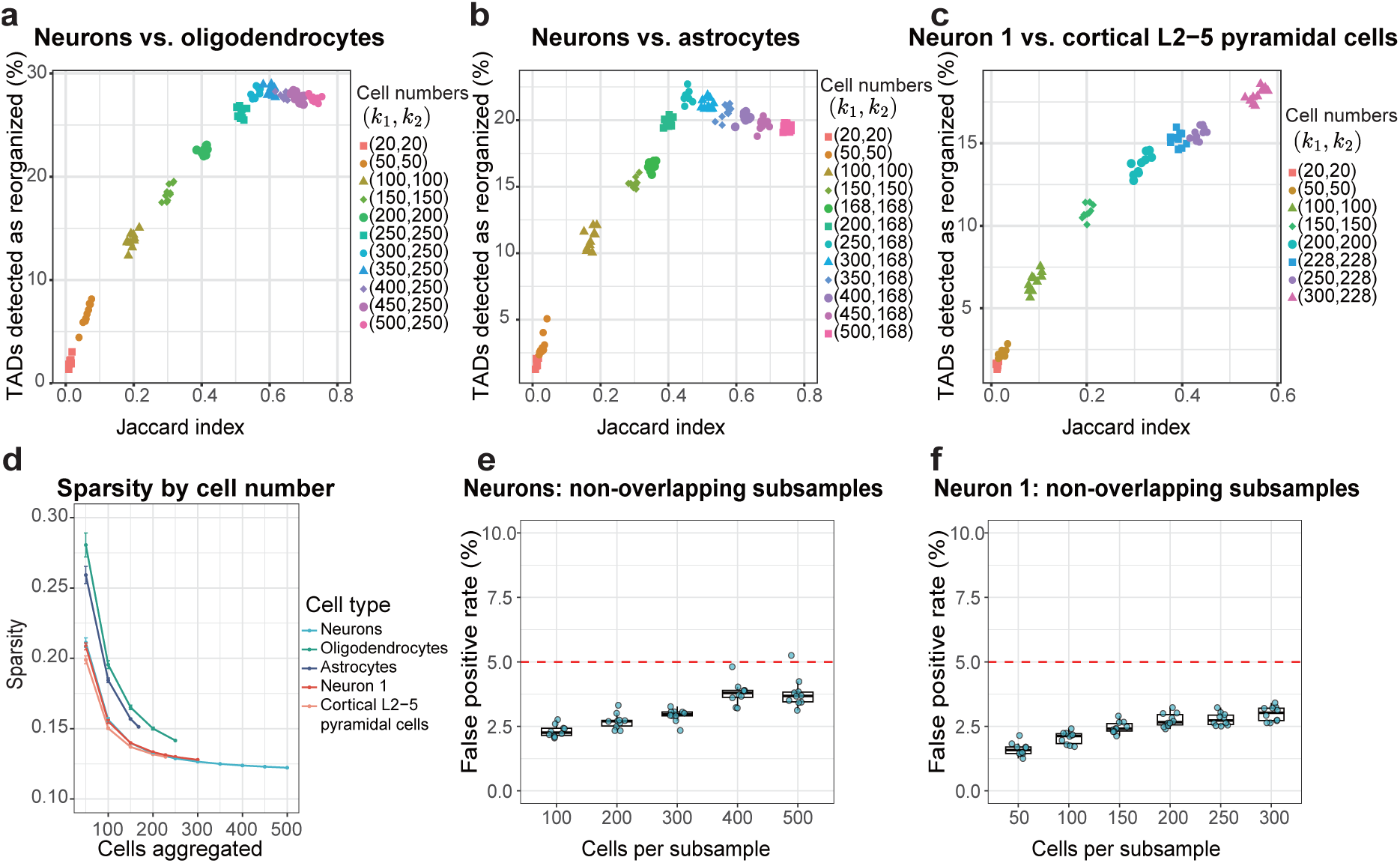
Effects of cell number on sparsity, detection reproducibility and false positive control of DiffDomain-Spectrum. **a–c**, Detection proportion and reproducibility across cell-number settings for comparisons between **a**, neurons and oligodendrocytes; **b**, neurons and astrocytes; and **c**, neuron 1 and cortical L2–5 pyramidal cells. For each comparison, cells were randomly subsampled without replacement, and their raw contacts were pooled to construct aggregated scHi-C contact maps at 25 kb resolution. Each point represents one subsampling repeat. Colors and symbols denote the cell-number pair (*k*_1_, *k*_2_), where *k*_1_ and *k*_2_ are the numbers of cells aggregated from the reference and comparison cell types, respectively. The y-axis shows the proportion of reference TADs detected as reorganized, and the x-axis shows the Jaccard index relative to the corresponding call set obtained using all available cells. Neurons-defined TADs were used as the reference set in **a** and **b**, and neuron 1-defined TADs were used in **c**. **d**, Sparsity of aggregated scHi-C contact maps constructed from increasing numbers of neurons, oligodendrocytes, astrocytes, neuron 1 cells and cortical L2– 5 pyramidal cells. Sparsity was defined as the proportion of zero-valued entries across contact submatrices corresponding to the tested TADs. Points show the mean across subsampling repeats, and error bars indicate the standard deviation across subsampling repeats. **e,f**, Empirical FPRs obtained from pairs of non-overlapping cell sets sampled without replacement from the same cell type and aggregated separately to construct paired aggregated scHi-C contact maps for **e**, neurons and **f**, neuron 1. The x-axis shows the number of cells in each subsample. Points represent individual subsampling repeats; boxes show the median and interquartile range, and whiskers extend to 1.5*×* the interquartile range. The red dashed line denotes the nominal 5% FPR level.

Increasing the cell number in only one condition produced substantially smaller gains in reproducibility. For example, after aggregating 250 neurons and 250 oligodendrocytes, increasing the number of neurons from 250 to 500 while fixing the number of oligodendrocytes at its maximum available value of 250 reduced the mean sparsity of the neurons contact map from 0.129 to 0.122, but produced only a limited increase in the Jaccard index and a slight decrease in the proportion of TADs detected as reorganized (Fig. 4a,d). Similarly limited improvement was observed when the number of neurons was increased while the number of astrocytes remained fixed at its maximum available value (Fig. 4b,d). These results indicate that increasing cell numbers in both conditions improves reproducibility more effectively than increasing the cell number in only one condition.

To determine whether varying cell numbers affected false positive control, we sampled two non-overlapping cell sets without replacement from the same biological condition and aggregated each set separately to construct a pair of replicate aggregated scHi-C contact maps. Across all tested cell-number settings for neurons and neuron 1, empirical FPRs remained below the nominal 5% level, with only a small number of subsampling repeats yielding FPRs that slightly exceeded this threshold (Fig. 4e,f). Thus, increasing the number of aggregated cells improved reproducibility without systematically inflating the empirical FPR.

We next evaluated the effect of genomic resolution using LiMCA scHi-C data from the human cell lines GM12878, K562, BJ and eHAP [51]. DiffDomain-Spectrum was applied to six pairwise contrasts among these cell lines, with each contrast evaluated in both reference directions, yielding 12 reference-TAD comparisons at both 25 kb and 50 kb resolutions. For each direction, reference TADs called from the corresponding bulk Hi-C map were used as a common genomic interval set at both resolutions. Increasing the bin size from 25 kb to 50 kb increased the proportion of TADs detected as reorganized in all comparisons, with a mean absolute increase of 11.5 percentage points (Fig. 5a,b). This increase is consistent with the lower sparsity and greater contact coverage at the coarser 50 kb resolution.

**Figure 5:**
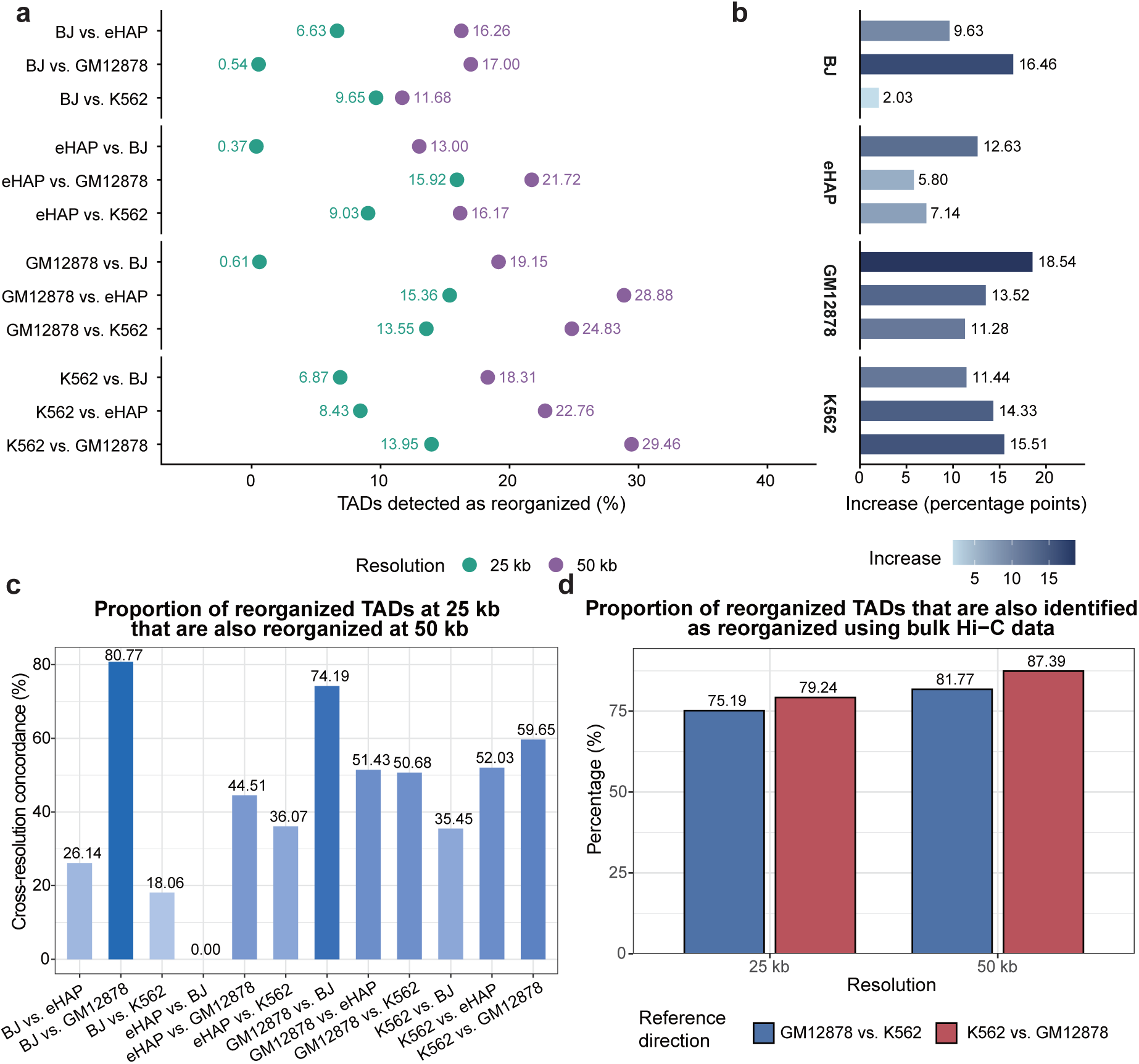
Effects of genomic resolution and concordance with matched bulk Hi-C data for DiffDomain-Spectrum. **a**, Proportion of reference TADs detected as reorganized from LiMCA aggregated scHi-C contact maps at 25 kb and 50 kb resolution across 12 reference-TAD comparisons among GM12878, K562, BJ and eHAP. The four cell lines form six pairwise contrasts, each evaluated in both reference directions. For each direction, reference TADs called from the corresponding bulk Hi-C map were used as a common genomic interval set at both resolutions. **b**, Absolute increase in the proportion of TADs detected as reorganized when the bin size was increased from 25 kb to 50 kb. Values were calculated as the proportion at 50 kb minus that at 25 kb and are expressed in percentage points. **c**, Cross-resolution concordance of DiffDomain-Spectrum calls, defined among reference intervals yielding valid results at both resolutions as the proportion of TADs detected as reorganized at 25 kb that were also detected as reorganized at 50 kb. Concordance was at least 50% in six of the 12 reference-TAD comparisons. Results for the alternative bulk callers are shown in Supplementary Fig. 3. **d**, Concordance between reorganized TADs detected from aggregated scHi-C contact maps and matched bulk in situ Hi-C data for GM12878 and K562. DiffDomain-Spectrum was applied to both data types using the same bulk-Hi-C-derived reference TAD intervals. For each reference direction and resolution, bars show the proportion of TADs detected as reorganized from aggregated scHi-C contact maps that were also detected as reorganized from matched bulk Hi-C data.

Among reference intervals yielding valid results at both resolutions, cross-resolution concordance was calculated as the proportion of TADs detected as reorganized at 25 kb that were also detected as reorganized at 50 kb. For DiffDomain-Spectrum, this proportion was at least 50% in six of the 12 reference-TAD comparisons (Fig. 5c). The corresponding numbers were nine comparisons for DiffDomain, eight for HiCcompare and all 12 for HiCDC+, whereas none of the remaining alternative bulk callers exceeded 50% overlap in any comparison (Supplementary Fig. 3). These higher overlap frequencies do not necessarily indicate superior overall performance. The higher overlap of DiffDomain may partly reflect its conservative detection behavior and smaller call sets. HiCcompare and HiCDC+ were adapted here by summing intra-TAD interaction counts and testing differences in total counts, a summary statistic expected to be comparatively insensitive to changes in binning resolution. Their overlap frequencies must also be interpreted together with their benchmarking results: HiCcompare showed conservative detection, whereas HiCDC+ exhibited markedly inflated FPRs. Overall, DiffDomain-Spectrum retained moderate cross-resolution concordance while maintaining a more favorable balance between false positive control and detection sensitivity.

Finally, we assessed concordance with matched bulk data by applying DiffDomain-Spectrum separately to aggregated scHi-C and bulk in situ Hi-C contact maps for GM12878 and K562 using the same bulk-Hi-C-derived reference TAD intervals [4]. In both reference directions and at both resolutions, more than 75% of reorganized TADs detected from aggregated scHi-C contact maps were also detected from matched bulk Hi-C data. The overlap reached 87.39% for the K562-versus-GM12878 comparison at 50 kb resolution (Fig. 5d). Thus, most reorganized TADs detected from the aggregated scHi-C contact maps were concordant with those detected from matched bulk Hi-C data.

Together, these analyses show that reproducibility increases with the number of aggregated cells while empirical FPR remains controlled. DiffDomain-Spectrum also retains moderate cross-resolution concordance and substantial agreement with matched bulk Hi-C data, supporting robust application across cell numbers and genomic resolutions.

### Reorganized TADs show subtype-consistent CTCF changes and are enriched for differentially expressed genes

To assess the biological relevance of reorganized TADs, we first examined changes in CTCF binding at TAD boundaries. CTCF is an architectural protein that contributes to chromatin looping and TAD boundary insulation, and changes in boundary-associated CTCF occupancy accompany TAD reorganization [2, 52]. Previous analyses of bulk Hi-C data using DiffDomain found that strength-change-up TADs with increased contact frequencies tended to show more CTCF binding sites at retained boundaries, whereas boundaries lost in loss, zoom and merge subtypes tended to show fewer CTCF sites [26]. These observations motivate directional expectations at the group level but do not imply a deterministic relationship between intra-domain contact strength and CTCF occupancy. We therefore asked whether reorganized TADs detected by DiffDomain-Spectrum showed boundary-associated CTCF changes broadly concordant with their structural subtypes. Reorganized TADs detected by DiffDomain-Spectrum were assigned to structural subtypes following the DiffDomain framework, as described in Methods (Supplementary Fig. 4a). We quantified CTCF peak counts at subtype-associated boundaries across six pairwise LiMCA contrasts evaluated in both reference directions, yielding 12 reference-TAD comparisons. For each comparison, CTCF peak counts were measured using condition-2 data and compared with counts at condition-2 TAD boundaries that did not overlap reorganized condition-1 TADs.

In the GM12878-versus-K562 comparison, DiffDomain-Spectrum detected 778 reorganized TADs distributed across the structural subtypes (Supplementary Fig. 4b). Relative to the “other boundaries” control group, *strength-change up* boundaries had significantly higher CTCF peak counts, whereas boundaries associated with *strength-change down*, *zoom*, *merge* and *complex* TADs had significantly lower counts (one-sided Mann–Whitney U tests; Supplementary Fig. 4c). Across all 12 reference-TAD comparisons, subtype composition varied among cell-line contrasts (Supplementary Fig. 4d), and multiple subtype–comparison combinations showed significant increases or decreases in CTCF peak counts relative to the control group. The directions of these changes were broadly consistent with boundary retention or loss and with the subtype-associated CTCF patterns previously observed in bulk Hi-C data (Supplementary Fig. 4e), supporting the structural relevance of the reorganized TADs detected by DiffDomain-Spectrum.

We next integrated matched scRNA-seq data to examine transcriptional associations with TAD reorganization. Because scRNA-seq and scHi-C were co-assayed in the same individual cells, the two modalities could be integrated within directly matched cell populations without cross-sample matching. DEGs were significantly enriched among expressed genes located within reorganized TADs in both the GM12878-versus-K562 and GM12878-versus-eHAP comparisons (hypergeometric test, *P* = 1.05 *×* 10*^−^*^7^ and *P* = 2.56 *×* 10*^−^*^6^, respectively; Supplementary Fig. 5a,b). These overlaps included 599 DEGs in the GM12878-versus-K562 comparison and 604 DEGs in the GM12878-versus-eHAP comparison. Gene Ontology enrichment analysis of the overlapping genes identified terms related to B-cell activation and differentiation, lymphocyte and leukocyte proliferation, cell adhesion, immune receptor activity and cytoskeletal organization [53] (Supplementary Fig. 5c,d).

Together, the subtype-consistent CTCF changes and significant DEG enrichment support the biological relevance of the reorganized TADs detected by DiffDomain-Spectrum.

### Ablation analysis of preprocessing and goodness-of-fit testing

We assessed three preprocessing steps and alternative goodness-of-fit tests in DiffDomain-Spectrum. Using scNanoHi-C2 data from three GM12878 biological replicates and K562 at 50 kb resolution, we individually removed three preprocessing operations applied to the log-scale TAD-level difference matrix: clipping finite entries to [−3, 3], replacing all-missing-offset with standard-normal random draws, and replacing +∞ and −∞with distance-specific finite extrema.

Removing extreme-value clipping increased FPRs to 8.10%, 10.30% and 11.38% across the three GM12878 replicate comparisons and increased the detection proportion from 18.80% to 22.84% in GM12878 versus K562 (Supplementary Fig. 6a,b), showing that increased detection yield came at the cost of false positive control. Omitting all-missing-offset replacement had little effect on FPR or detection proportion but substantially reduced the number of TADs retained for analysis (Supplementary Fig. 6a–c). Omitting distance-specific replacement of infinite entries had little effect on FPR and modestly reduced the detection proportion to 18.03% (Supplementary Fig. 6b).

We next compared goodness-of-fit tests within the same random matrix theory framework at 25 kb resolution. Detection proportions were evaluated using LiMCA GM12878-versus-K562 and GM12878-versus-eHAP comparisons, and false positive control using dscHi-C mESC biological-replicate and replicate-versus-subsample controls. The full-spectrum goodness-of-fit test used by DiffDomain-Spectrum, denoted *Z_A_*, was compared with Pearson’s chi-squared, Anderson–Darling (AD) [54], Kolmogorov– Smirnov (KS) [55], Cramér–von Mises (CvM) [56, 57], *Z_K_* and *Z_C_* [38].

*Z_A_* detected 13.55% and 15.36% of reference TADs as reorganized while maintaining FPRs of 0.08% and 0.30% (Supplementary Fig. 6d,e). Pearson’s chi-squared test increased detection proportions to 29.47% and 30.79% but yielded markedly inflated FPRs of 31.51% and 31.72%. A similar trade-off was observed for *Z_K_*. AD, KS, CvM and *Z_C_* maintained low FPRs but detected no reorganized TADs in either biological comparison. *Z_A_* therefore achieved the highest detection proportion among tests maintaining FPR below 5% in both controls.

Together, extreme-value clipping contributed to false positive control, all-missing-offset replacement increased the number of analyzable TADs, and *Z_A_*provided the most favorable balance between detection sensitivity and false positive control.

## Discussion

We present DiffDomain-Spectrum, a statistically principled framework for identifying structurally reorganized TADs between conditions or cell types from sparse aggregated scHi-C contact maps. The method tests normalized TAD-level difference matrices derived from aggregated raw scHi-C contact maps and leverages the full eigenvalue spectrum to detect domain-level contact changes, without separately normalizing sparse maps or enhancing individual scHi-C contact maps. Accordingly, DiffDomain-Spectrum avoids enhancing individual scHi-C contact maps and performs inference from transformed summaries of observed contacts in the aggregated maps. Across multiple scHi-C platforms, DiffDomain-Spectrum achieved a favorable balance between false positive control and detection sensitivity relative to alternative bulk callers, while consistently detecting a higher proportion of reference TADs as reorganized than the boundary-focused single-cell method scHiCluster. Detected reorganized TADs showed coherent aggregate contact patterns, were associated with changes in CTCF binding and were enriched for differentially expressed genes, supporting their biological relevance.

DiffDomain-Spectrum is designed for aggregated scHi-C contact maps that remain substantially sparse at biologically informative resolutions such as 25 kb despite gains in coverage from aggregating contacts across cells. In this setting, DiffDomain-Spectrum complements existing bulk Hi-C callers, including DiffDomain, by extending comparative domain-level analysis to sparse aggregated scHi-C contact maps, for which these callers were not specifically developed. Relative to the boundary-focused single-cell method scHiCluster, DiffDomain-Spectrum targets domain-level contact-pattern reorganization in aggregated maps rather than differential TAD boundary occurrence after enhancing individual scHi-C contact maps. As scHi-C and single-cell multiomic profiling increasingly encompass heterogeneous tissues, developmental trajectories, aging and disease [16, 58–60], DiffDomain-Spectrum is positioned to support comparative analysis of domain organization across these increasingly diverse biological settings.

Aggregation supports stable domain-level testing between conditions or cell types but collapses the distribution of contact patterns across individual cells. DiffDomain-Spectrum therefore identifies population-level TAD reorganization but does not determine whether a detected change is broadly shared across cells or driven by a subset of cells. Extending DiffDomain-Spectrum to model and compare cell-level distributions of domain contact patterns could reveal, for example, whether detected reorganization is broadly shared, varies in magnitude, reflects shifts among structural states or is concentrated in specific subpopulations.

Together, these results establish DiffDomain-Spectrum as a statistically principled framework for identifying structurally reorganized TADs from sparse aggregated scHi-C contact maps. Because it requires neither separate normalization of sparse maps nor enhancement of individual scHi-C contact maps, the framework provides a practical basis for comparing TAD reorganization across an expanding range of cell types, developmental trajectories, perturbations and disease states profiled by scHi-C.

## Methods

### DiffDomain-Spectrum framework

#### Construction of aggregated scHi-C contact maps and reference TADs

For each condition or cell type, observed contacts from all retained cells were aggregated by summing contact counts at identical pairs of genomic bins. For datasets provided as individual-cell contact-pair files, files from cells in the same group were combined before conversion to binned contact maps. No per-cell weighting, normalization, smoothing or enhancement was applied before aggregation. Subsequent TAD-level analyses used chromosome-specific cis contact matrices. Contact maps at 25 and 50 kb resolution were constructed independently from the underlying contacts rather than by rebinning maps generated at another resolution. Unless otherwise specified, analyses were performed at 25 kb resolution. The scNanoHi-C2 data were analyzed at 50 kb resolution, and the LiMCA data were analyzed at both 25 and 50 kb resolution.

DiffDomain-Spectrum accepts a predefined reference TAD set and does not itself call TADs. In the default workflow, reference TADs were called from the aggregated scHi-C contact map for condition 1 using the insulation-score method implemented in FAN-C v0.9.28 [61]. Boundaries were identified from the 200 kb insulation-score track using a minimum boundary score of 0.3, and TAD intervals were defined between consecutive boundaries independently within each chromosome. TADs were also called from the condition-2 aggregated contact map using the same method and parameters solely for structural-subtype classification and did not determine the genomic intervals tested by DiffDomain-Spectrum. When both directions of a comparison were analyzed, reference TADs were called separately from the map designated as condition 1 in each direction. For comparisons with the boundary-focused single-cell method scHiCluster, TopDom [62] was used instead of the insulation-score method, and the condition-1 TopDom TADs provided the common reference set for evaluating the two methods. For the LiMCA cross-resolution and matched-bulk concordance analyses, reference TADs called from the corresponding bulk Hi-C maps were used to provide a common interval set across aggregated scHi-C and bulk Hi-C analyses.

#### Construction and preprocessing of TAD-level difference matrices

For each condition-1 reference TAD, square symmetric contact-count matrices spanning the reference interval were extracted from the aggregated contact maps for conditions 1 and 2 and denoted by *M*_1_ and *M*_2_, respectively. Diagonal and near-diagonal entries were retained. Entries with zero contact counts were treated as missing. For each genomic bin, the proportion of missing entries was calculated separately in *M*_1_ and *M*_2_, including the diagonal entry. A bin was removed from both matrices when this proportion was at least 50% in either condition. This filtering was applied once, with identical rows and columns removed from *M*_1_ and *M*_2_. The number of retained bins was denoted by *N*, and TADs retaining fewer than 10 bins were excluded from subsequent testing.

A log-scale TAD-level difference matrix was constructed element-wise as

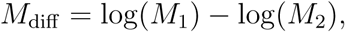

where log denotes the natural logarithm and no pseudocount was added. Entries of *M*_diff_ were then stratified by their matrix diagonal offset *k* =|*i − j|*, with *k* = 0*,…, N −* 1. For each offset containing finite entries, the median, maximum, mean and standard deviation were calculated from the finite upper-triangular entries before replacement. Offset-specific standard deviations that were zero or undefined were set to one. Missing entries were replaced with the offset-specific median, whereas +∞ and −∞ were replaced with the offset-specific maximum finite value and its negative, respectively.

After these replacements, *M*_diff_ was standardized within each matrix diagonal offset as

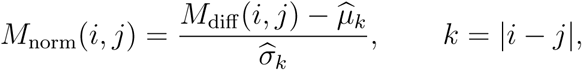

where *μ̂_k_* and *σ̂_k_* denote the mean and standard deviation calculated from the original finite entries at offset *k*. For offsets at which all entries were missing, standard-normal random values were assigned directly to the offsets, with the same draw assigned to each symmetric pair to guarantee the symmetry of *M*_norm_. Finally, *M*_norm_ was clipped element-wise to the interval [−3, 3].

#### Spectral null model based on random matrix theory

For each reference TAD retaining at least 10 bins after sparse-bin removal, the normalized difference matrix was scaled by the square root of its dimension,

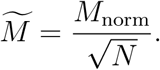

Because entries of *M*_norm_ were centered and scaled within matrix diagonal offsets, division by 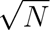 reduces their variances to order 1*/N* and prevents the eigenvalue spectrum from increasing in scale with matrix dimension.

Following the random matrix theory formulation of DiffDomain [26], the spectral null hypothesis of no structural reorganization is motivated by generalized Wigner matrices. A generalized Wigner matrix is symmetric, with independent, mean-zero upper-triangular entries whose variances are of order 1*/N*. The construction of *M̃* produces a symmetric matrix with approximately centered and variance-standardized entries, but the independence assumption is not satisfied by aggregated scHi-C contact maps. Contact frequencies among nearby chromosome-bin pairs can be strongly correlated, inducing local dependence among entries of *M̃*. We therefore use the generalized Wigner model as an approximate spectral null rather than assuming that *M̃* is an exact generalized Wigner matrix. As shown previously for DiffDomain, this local dependence does not prevent normalized Hi-C difference matrices from exhibiting the empirical spectral behavior predicted by random matrix theory [26]. DiffDomain-Spectrum extends this formulation by evaluating the full eigenvalue spectrum rather than only its largest eigenvalue.

Because M̃ is real and symmetric, it has *N* real eigenvalues, which were ordered as

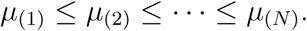

All *N* eigenvalues were retained. Their empirical cumulative distribution function was defined as

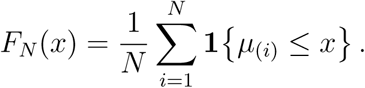

Under the spectral null model, the empirical spectral distribution *F_N_* is expected to approximate the standard semicircle law. Its density is

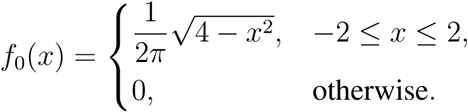

Let *F*_0_ denote the corresponding cumulative distribution function. Systematic departure of *F_N_*from *F*_0_ provides evidence of structural reorganization. This departure was quantified using the goodness-of-fit procedure described below.

#### Statistical inference

Departure of the observed eigenvalue spectrum from the semicircle law was quantified using the likelihood-ratio goodness-of-fit statistic *Z_A_*[38]. The null hypothesis was that the empirical eigenvalue spectrum was compatible with the semicircle-law spectral null, whereas the alternative was a systematic departure from this distribution. As an omnibus statistic, *Z_A_* incorporates all ordered eigenvalues, enabling detection of distributed departures across the full spectrum rather than reducing the matrix to a single extreme eigenvalue.

For the ordered eigenvalues *µ*_(1)_*,…, µ*_(_*_N_*_)_, let *u_i_* = *F*_0_(*µ*_(_*_i_*_)_), with cumulative probabilities truncated to [10*^−^*^10^, 1 − 10*^−^*^10^] to avoid evaluating log(0). The observed statistic was calculated as

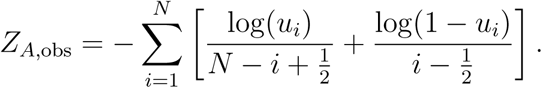

All eigenvalues contribute additively to *Z_A_*, but their contributions are rank dependent rather than equal. The negative of the bracketed term at rank *i* gives the contribution of *µ*_(_*_i_*_)_ and depends jointly on its semicircle-law cumulative probability and its rank within the ordered spectrum. The two rank-dependent denominators are exchanged between mirrored ranks *i* and *N* + 1 − *i*, providing symmetric treatment of lower- and upper-tail departures. For each rank, the contribution is minimized at the continuity-corrected null rank position 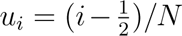 and increases as *u_i_* departs from this position. Consequently, moderate departures across the spectral bulk can accumulate, while substantial departures near either spectral tail can also contribute strongly.

Because the classical finite-sample null distribution of *Z_A_* is not available in a convenient closed form and the ordered eigenvalues do not constitute an independent and identically distributed sample, classical asymptotic calibration was not used. A raw *P*-value was instead estimated separately for each TAD using *B* = 10,000 Monte Carlo replicates. TAD-specific calibration accounts for variation in both the spectrum size *N* and the observed spectral range among reference TADs. Null eigenvalue sets were sampled directly from the theoretical semicircle distribution rather than generated by subsampling cells or contacts and reconstructing aggregated scHi-C contact maps. This design also avoids repeated construction and eigendecomposition of simulated matrices, thereby reducing computational time and memory requirements.

For each replicate, *N* values were sampled from the standard semicircle distribution conditional on falling within

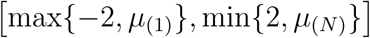

and were then ordered. The corresponding simulated statistic, *Z_A,r_*, was calculated using the same cumulative-probability truncation and formula as *Z_A,_*_obs_. Larger values of *Z_A_* indicate greater departure from the semicircle-law null, and the raw *P*-value was therefore calculated from the upper tail of the simulated statistics as

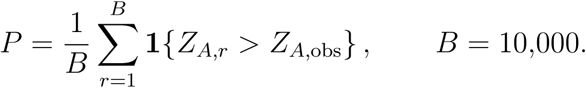

The Monte Carlo procedure is an approximate, method-specific calibration rather than an exact simulation of finite-dimensional random-matrix spectra. It samples eigenvalues independently from the semi-circle distribution and conditions the sampling range on the observed minimum and maximum eigenvalues. Consequently, it does not reproduce the dependence and eigenvalue repulsion among eigenvalues of a finite-dimensional Wigner matrix, and the calibration is data dependent. This approximation was adopted to avoid repeated generation and eigendecomposition of random matrices. Its practical adequacy was evaluated through empirical false positive control across ten control comparisons spanning three scHi-C platforms.

Once the aggregated maps have been constructed, calibration for each TAD depends on the matrix dimension and observed spectral range rather than on the number of contributing cells. TAD-level calibrations can be computed separately before joint Benjamini–Hochberg correction, making this stage naturally parallelizable.

For each directed comparison between two conditions or cell types, Benjamini–Hochberg correction was applied jointly to all valid raw *P*-values. TADs for which matrix construction, eigendecomposition or statistical testing did not yield a valid *P*-value were excluded from multiple-testing correction. A TAD was identified as structurally reorganized when its adjusted *P*-value was less than or equal to 0.05.

### Benchmarking and method evaluation

#### Alternative bulk callers

DiffDomain-Spectrum was compared with the alternative bulk callers DiffDomain, DiffGR, TADCompare, DiffTAD-parametric, DiffTAD-permutation, HiCcompare and HiCDC+ [26, 41–45]. All methods were applied to the same aggregated scHi-C contact maps and, where supported, the same condition-1 reference TAD set.

DiffDomain was applied to aggregated raw contact maps without additional contact-map normalization, using a minimum of ten retained bins per TAD. DiffGR was run using resolution-specific smoothing windows of 11 bins at 25 kb and 5 bins at 50 kb, Knight–Ruiz normalization, 100 permutations, the default stratum-adjusted correlation cutoff of 0.85 and its accelerated implementation. TADCompare was supplied with the condition-1 reference TAD set for both input maps. DiffTAD was run using the authors’ open-source implementation in its parametric and permutation modes, referred to as DiffTAD-parametric and DiffTAD-permutation, respectively.

HiCcompare and HiCDC+ were adapted for TAD-level analysis by representing each reference TAD as a single interaction feature whose coordinates were defined by the TAD interval and whose interaction strength was the sum of contacts within that interval. For HiCDC+, replicate-level intra-TAD contact counts were analyzed using its DESeq2-based differential-testing procedure.

For each directed comparison, valid method-specific *P*-values were adjusted using the Benjamini– Hochberg procedure. Reference TADs with adjusted *P ≤* 0.05 were considered reorganized.

#### Comparison with scHiCluster

DiffDomain-Spectrum was compared with the boundary-focused single-cell method scHiCluster using HiRES data from six developing mouse embryo cell types [20, 25]. All 30 directed pairwise comparisons among the six cell types were evaluated at 25 kb resolution.

Individual scHi-C contact maps were enhanced using the scHiCluster workflow with a two-bin convolutional padding window, square-root vanilla-coverage normalization, a random-walk restart probability of 0.5 and a maximum genomic distance of 2 Mb. TopDom was then applied to each enhanced individual contact map using a ten-bin window to identify TAD boundaries [62]. Differential boundary occurrence between cell populations was evaluated using the scHiCluster procedure with multiple-testing correction. For each directed comparison, TopDom TADs called from the condition-1 aggregated raw scHi-C contact map provided the common reference TAD set. Because differential boundaries identified from enhanced individual maps did not necessarily coincide exactly with the boundaries of these reference TADs, a reference TAD was considered reorganized by scHiCluster when at least one significant differential boundary fell within its genomic interval. DiffDomain-Spectrum and scHiCluster were then compared using the same reference TAD set.

#### Classification of reorganized TADs

Reorganized TADs were assigned to structural subtypes by comparing the condition-1 reference TAD set with TADs called from the condition-2 aggregated contact map, following the classification framework introduced for DiffDomain [26]. Two boundaries were considered matched when their genomic separation did not exceed the larger of 30 kb and 10% of the condition-1 TAD length. TADs connected through reciprocal interval overlap were considered jointly, with connected regions spanning more than 1 Mb excluded from further expansion.

A one-to-one correspondence with both boundaries matched was classified as *strength-change*. A one-to-one overlap without matched boundaries was classified as *zoom*. A condition-1 TAD corresponding to multiple condition-2 TADs was classified as *split*, whereas multiple condition-1 TADs corresponding to one condition-2 TAD were classified as *merge*. A condition-1 TAD without an overlapping condition-2 TAD was classified as *loss*. Remaining many-to-many or otherwise unresolved relationships were classified as *complex*.

Strength-change TADs were further divided according to their direction of contact-intensity change. Let *m*_1_ and *m*_2_ denote the median intra-TAD contact frequencies for the matched TAD in conditions 1 and 2, respectively, and let *s*_1_ and *s*_2_ denote the sums of intra-TAD contact frequencies across all TADs in the corresponding conditions. A strength-change TAD was classified as *strength-change up* when

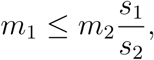

indicating greater contact intensity in condition 2 after global scaling, and as *strength-change down* otherwise.

#### Detection proportion and false positive rate

The detection proportion for a directed biological comparison was calculated as the number of condition-1 reference TADs detected as reorganized divided by the number of reference TADs yielding valid test results. Because no ground-truth set of reorganized TADs was available, this quantity was used to describe detection extent rather than statistical power.

Empirical false positive rates were evaluated using ten control comparisons between datasets representing the same biological condition. These comprised three biological-replicate comparisons from mouse cortex Dip-C data, four mESC dscHi-C controls consisting of one biological-replicate comparison, two replicate-versus-subsample comparisons and one biotin-versus-no-biotin comparison, and three pairwise comparisons among GM12878 scNanoHi-C2 biological replicates. All TADs detected as reorganized in these controls were treated as false positives, and the empirical false positive rate was calculated as their proportion among reference TADs with valid test results.

For the comparison with scHiCluster, directional overlap was calculated using the TAD-level calls derived from scHiCluster differential boundaries as the denominator. Specifically, it was the proportion of reference TADs considered reorganized by scHiCluster that were also detected as reorganized by DiffDomain-Spectrum.

#### Cell-number subsampling and reproducibility analysis

The effect of cell number was evaluated using mouse cortex Dip-C data at 25 kb resolution. For each cell type and cell-number setting, cells were randomly sampled without replacement in ten independent subsampling repeats, and their raw contacts were summed to construct aggregated scHi-C contact maps. Balanced settings used equal cell numbers in both conditions. In unbalanced settings, the number of cells in one condition was increased while the other condition was fixed at its maximum available cell number.

For each subsampling repeat, detection proportion was calculated among reference TADs with valid test results. Reproducibility was quantified using the Jaccard index between the subsample-derived reorganized-TAD set and the corresponding set obtained using all available cells,

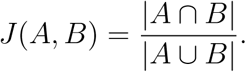

Sparsity was calculated as the proportion of zero-valued entries within the contact submatrices corresponding to the tested TADs.

To assess false positive control at different cell numbers, two non-overlapping cell sets were sampled without replacement from the same cell type and aggregated separately. The empirical false positive rate was calculated for each paired subsampling repeat using the same definition as for the control comparisons above.

#### Cross-resolution and matched bulk Hi-C concordance

Cross-resolution concordance was evaluated using LiMCA data from GM12878, K562, BJ and eHAP. The six unordered cell-line contrasts were evaluated in both reference directions, yielding 12 reference-TAD comparisons at 25 kb and 50 kb resolutions. For each direction, reference TADs called from the corresponding bulk Hi-C map were used as a common genomic interval set at both resolutions. Among reference intervals yielding valid results at both resolutions, cross-resolution concordance was defined as the proportion of TADs detected as reorganized at 25 kb that were also detected as reorganized at 50 kb.

Concordance with matched bulk Hi-C data was evaluated for GM12878 and K562 in both reference directions at 25 kb and 50 kb resolution. DiffDomain-Spectrum was applied separately to aggregated scHi-C and matched bulk in situ Hi-C contact maps using the same bulk-Hi-C-derived reference TAD intervals. Concordance was calculated as the proportion of TADs detected as reorganized from aggregated scHi-C contact maps that were also detected as reorganized from matched bulk Hi-C data.

### Aggregated domain analysis

Aggregated domain analysis was performed using `coolpuppy` on 25 kb aggregated scHi-C contact maps. Expected cis-contact frequencies were estimated using `cooltools`, retaining the main and near-diagonal entries. TAD intervals and flanking regions of one TAD length on each side were rescaled to 99 × 99 matrices before aggregation.

For each directed comparison, the reference profile was generated from condition-1 reference TADs using the condition-1 aggregated contact map. Condition-2 profiles were generated separately for reorganized TADs assigned to each structural subtype. Contact enrichment was displayed on the log_10_ scale. Differences between a condition-2 subtype profile, *M*_2_, and the reference profile, *M*_1_, were displayed as

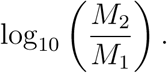

For comparisons among alternative bulk callers, aggregated domain analysis was restricted to methods detecting more than 5% of reference TADs as reorganized. The loss subtype was excluded when too few TADs were available for stable aggregation.

### Biological characterization of reorganized TADs

#### CTCF binding analysis

Boundary-associated CTCF binding was evaluated using publicly available CTCF ChIP–seq peak sets for the LiMCA cell lines. Reorganized TADs were assigned to the structural subtypes described above. For each subtype, retained, lost and gained boundaries were identified according to the correspondence between condition-1 and condition-2 TADs. CTCF peaks were counted within 30 kb on either side of each evaluated boundary using `bedtools intersect`.

All comparisons were performed using the CTCF data corresponding to condition 2. The control group, termed other boundaries, comprised condition-2 TAD boundaries that were not associated with condition-1 TADs detected as reorganized. CTCF peak-count distributions for each subtype-associated boundary group were compared with the other-boundaries group using one-sided Mann–Whitney *U* tests. Comparisons with *P <* 0.05 were displayed in the subtype-level summaries.

#### Differentially expressed gene enrichment

Matched scRNA-seq data from the LiMCA dataset were used to examine associations between TAD reorganization and gene-expression changes. Because scRNA-seq and scHi-C were co-assayed in the same individual cells, the two data types were analyzed in directly matched cell populations. Differential expression was assessed from the matched scRNA-seq count data using DESeq2 without aggregating counts across cells [63].

Genes with Benjamini–Hochberg-adjusted *P ≤* 0.05 and absolute log_2_ fold change of at least 2 were defined as differentially expressed genes. Human gene coordinates and HGNC symbols were obtained from Ensembl using `biomaRt`. A gene was assigned to a TAD when its genomic interval overlapped the TAD interval.

Enrichment of differentially expressed genes within reorganized TADs was evaluated using a one-sided hypergeometric test. The background comprised genes included in the corresponding differential-expression analysis that overlapped any tested reference TAD. The foreground comprised background genes overlapping TADs detected as reorganized. The test evaluated whether differentially expressed genes were overrepresented among genes located within reorganized TADs.

### Gene Ontology enrichment analysis

Gene Ontology enrichment analysis was performed on differentially expressed genes overlapping reorganized TADs using `clusterProfiler` and the human annotation database `org.Hs.eg.db`. HGNC gene symbols were used as input, and biological process, cellular component and molecular function ontologies were evaluated jointly. Enrichment *P*-values were adjusted using the Benjamini–Hochberg procedure. Terms with adjusted *P <* 0.05 and *q <* 0.2 were retained. Selected significant terms were displayed according to gene ratio, number of contributing genes and adjusted *P*-value.

### Ablation analyses

#### Ablation of difference-matrix preprocessing

The contributions of three difference-matrix preprocessing operations were evaluated using scNanoHi-C2 data at 50 kb resolution. The full DiffDomain-Spectrum procedure was compared with variants that individually omitted finite-value clipping to [−3,3], replacement of matrix diagonal offsets containing no finite values, or distance-specific replacement of positive and negative infinite entries. All remaining steps and statistical-testing procedures were unchanged.

False positive rates were evaluated across the three pairwise comparisons among GM12878 biological replicates. Detection proportion was evaluated in the GM12878-versus-K562 comparison. The effect of all-missing-offset replacement on analyzability was quantified as the relative increase in the number of reference TADs yielding valid test results compared with the corresponding analysis without this replacement.

### Comparison of goodness-of-fit tests

Alternative goodness-of-fit tests were evaluated within the same difference-matrix preprocessing and random-matrix framework at 25 kb resolution. The *Z_A_* statistic used by DiffDomain-Spectrum was compared with Pearson’s chi-squared, Anderson–Darling, Kolmogorov–Smirnov, Cramér–von Mises, *Z_K_*and *Z_C_* tests [38, 54–57]. Detection proportions were evaluated using the LiMCA GM12878-versus-K562 and GM12878-versus-eHAP comparisons. False positive rates were evaluated using an mESC dscHi-C biological-replicate comparison and a replicate-versus-subsample control. For each test and directed comparison, valid *P*-values were adjusted using the Benjamini–Hochberg procedure, and adjusted *P <* 0.05 was used to identify reorganized TADs.

## Data availability

All datasets used in this study are publicly accessible. The scHi-C datasets for mouse cortical cells (Dip-C) were downloaded from the Gene Expression Omnibus (GEO) under accession number GSE146397 [15]. The mouse brain cortex scHi-C dataset (dscHi-C) was obtained from GEO under accession number GSE285812 [39]. The scHi-C datasets for GM12878 (with biological replicates) and K562 generated using the scNanoHi-C2 approach were downloaded from GEO under accession number GSE269496 [40]. The scHi-C datasets for GM12878, K562, eHAP, and BJ generated using the LiMCA approach, together with the matched scRNA-seq data, were downloaded from GEO under accession number GSE240128 [51]. The matched bulk Hi-C datasets for GM12878 and K562 were downloaded from GEO under accession number GSE63525 [4]. Additionally, the scHi-C datasets for mouse embryo cells generated using the HiRES approach were downloaded from GEO under accession number GSE223917 [20]. The CTCF ChIP–seq peak sets for GM12878, K562, and BJ were obtained from the ENCODE (https://www.encodeproject.org/) project under accession numbers ENCSR000DZN, ENCSR000DWE and ENCSR000DQI, respectively. CTCF ChIP-seq data for HAP1 cells were obtained from GEO under accession number GSM4625027.

## Code availability

The software is published under the GNU GPL v3.0 license. The source code of DiffDomain-Spectrum is available at https://github.com/FocusPaka/DiffDomain-Spectrum.

## Acknowledgments

This work was supported by the National Natural Science Foundation of China grant 12271536 (D.T.), Natural Science Foundation of Guangdong Province of China grant 2024A1515012037 (Y.Z.), the National Natural Science Foundation of China grant 12371295 (Y.Z.), Shenzhen Science and Technology Program grant RCJC20221008092753082 (Y.Z.). Additionally, this work was supported by High-performance Computing Public Platform (Shenzhen Campus) of Sun Yat-sen University.

## Author contributions

D.T. conceived and supervised the study. J.Z., Y.Z. and D.T. developed the methodology. J.Z. and X.Z. developed the software. J.Z., Y.D., X.Z., H.Z., Y.Z. and D.T. performed the investigation and interpreted the results. J.Z. prepared the initial manuscript draft. H.Z. led substantive revision and rewriting of the manuscript, with D.T. providing overall scientific direction and editing. All authors reviewed and approved the final manuscript. Y.Z. and D.T. acquired funding.

## Supplementary Figures

**Supplementary Figure 1:**
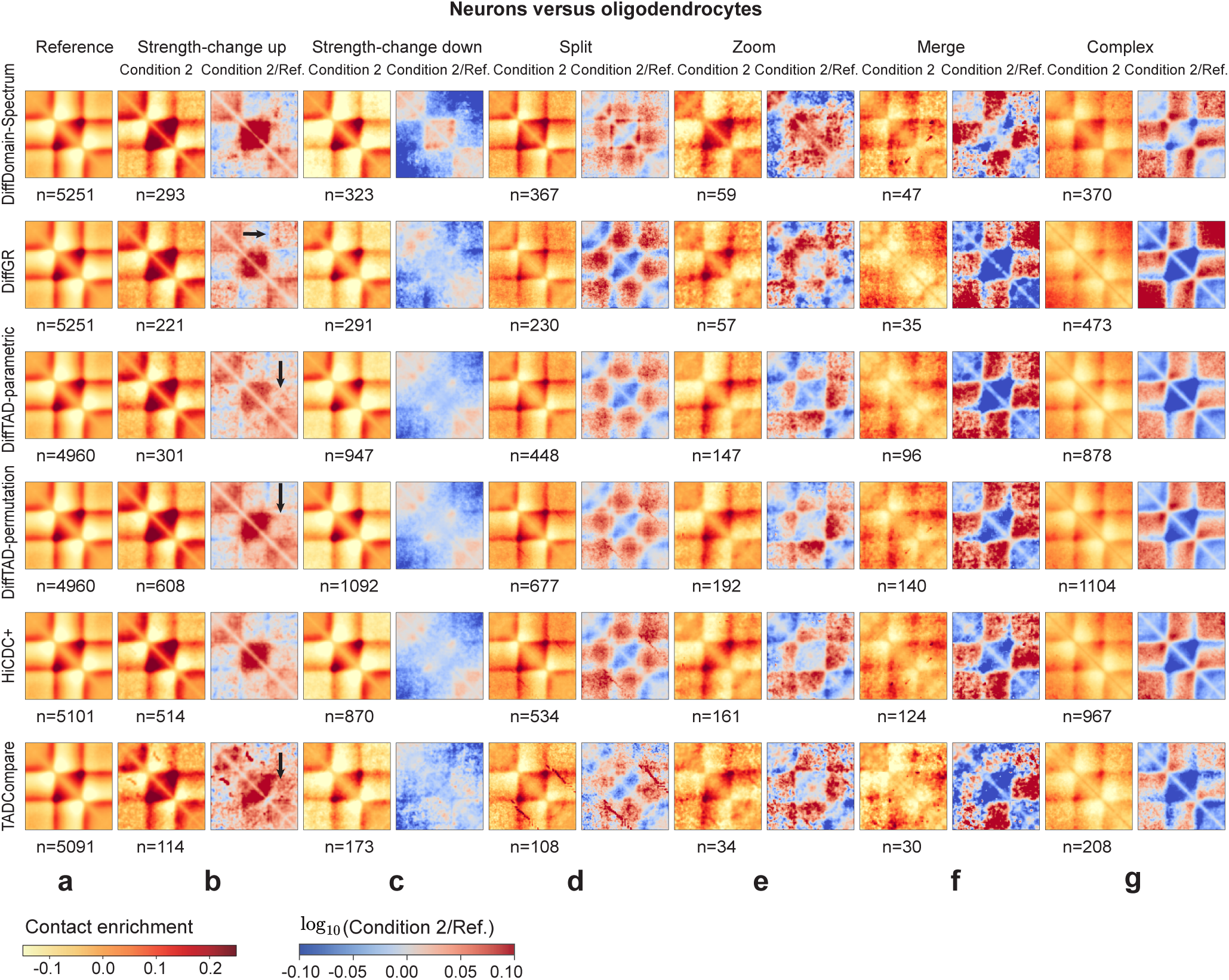
Aggregated domain analysis (ADA) plots summarizing reorganized TAD sub-types identified by six methods. Methods identifying more than 5% of TADs as reorganized are included. The number of aggregated TADs for each subtype is shown under the heatmap plot. *Loss* is excluded due to an insufficient number of reorganized TADs across six methods. **a** ADA plot summarizing all TADs in condition 1 based on aggregated scHi-C contact maps from neurons. The ADA plot is consistently used as a control in the subsequent ADA plots to highlight aggregated changes in the subtypes of reorganized TADs, and is referred to as condition 1 in subsequent ADA plots. **b** ADA plot summarizing aggregated changes in *strength-change up* TADs. Left: ADA plot for *strength-change up* TADs using aggregated scHi-C contact maps from oligodendrocyte (condition 2); Right: log_10_ fold-change between ADA matrices, where the numerator corresponds to the left panel and the denominator corresponds to panel **a**. For DiffGR, DiffTAD-parametric, DiffTAD-permutation, and TADCompare, interaction enrichment patterns spread beyond the expected TAD boundaries, making them less consistent with the definition of *strength-change up*. These regions are indicated by black arrows. **c** ADA plot summarizing aggregated changes in *strength-change down* TADs. Decreased interactions within the aggregated TAD region together with stable interactions around its boundaries support the definition of *strength-change down*. **d** ADA plot summarizing aggregated changes in *split* TADs. Decreased interactions within the aggregated TAD region, particularly around its center, are consistent with the definition of *split* TADs. **e** ADA plot summarizing aggregated changes in *zoom* TADs. Increased or decreased interactions within the aggregated TAD region and between it and adjacent upstream/downstream regions are typical of *zoom* TADs. **f** ADA plot summarizing aggregated changes in *merge* TADs. Increased interactions between the aggregated TAD region and its adjacent regions align with the definition of *merge* TADs. **g** ADA plot summarizing aggregated changes in *complex* TADs. Across the six methods (rows), distinct interaction changes within the aggregated TAD region and between the aggregated TAD and its adjacent regions are consistent with the interaction patterns expected for *complex* TADs. The ADA plots are generated using the Python package `coolpuppy` from aggregated scHi-C contact maps at 25 kb resolution.

**Supplementary Figure 2:**
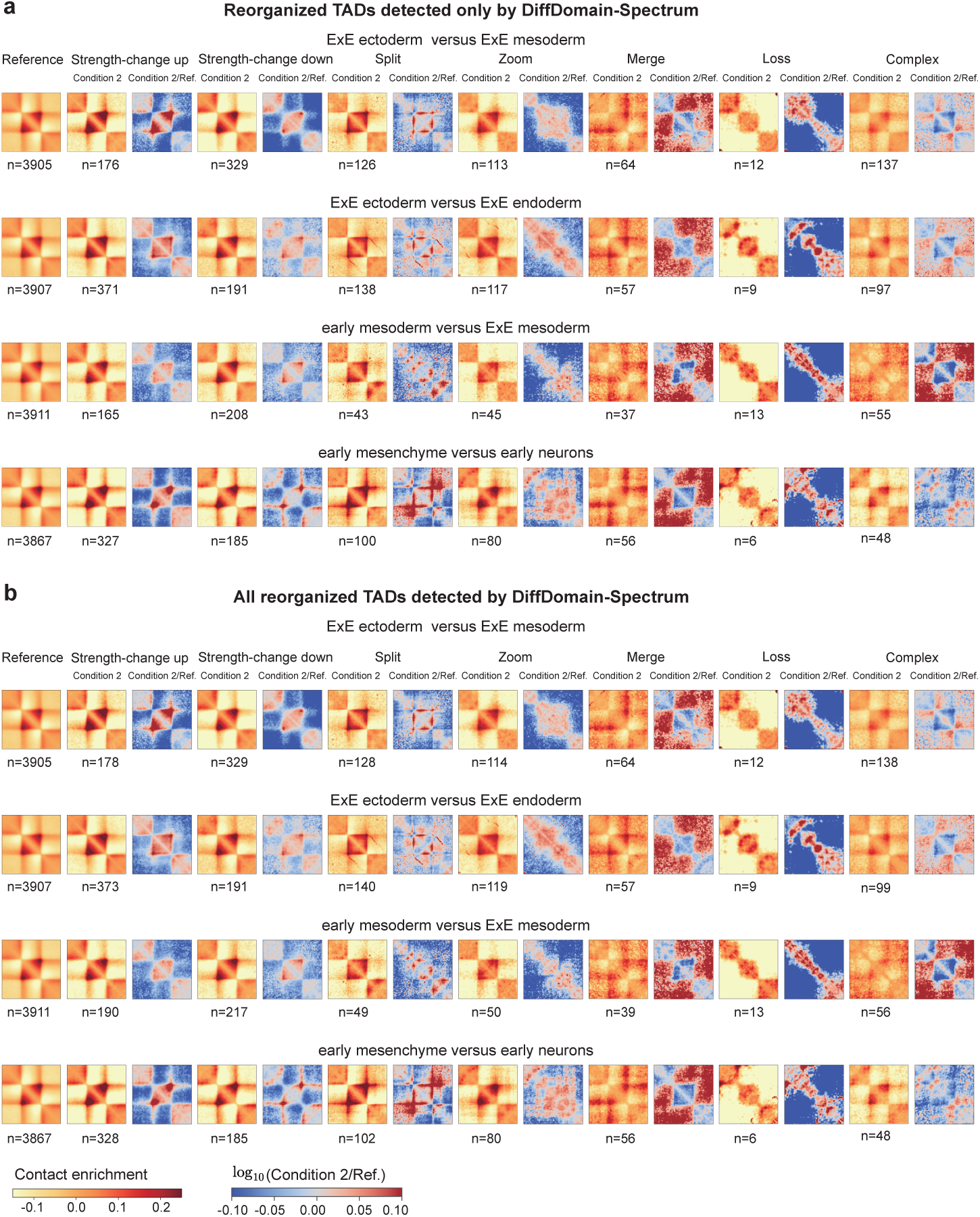
Subtype-level aggregated domain analysis of DiffDomain-Spectrum reorganized TADs in the comparison with scHiCluster. **a**, Aggregated domain analysis (ADA) of reorganized TADs detected only by DiffDomain-Spectrum across four representative directed comparisons. Reorganized TADs also classified as reorganized based on scHiCluster differential-boundary calls were excluded. Rows correspond to ExE ectoderm versus ExE mesoderm, ExE ectoderm versus ExE endoderm, early mesoderm versus ExE mesoderm and early mesenchyme versus early neurons. The Reference column shows the ADA profile of all condition-1 reference TADs using the condition-1 aggregated scHi-C contact map. For each structural subtype, the first map shows the ADA profile of reorganized TADs using the condition-2 aggregated scHi-C contact map, and the second shows log_10_(*M*_2_*/M*_1_), where *M*_1_ is the reference ADA matrix and *M*_2_ is the corresponding subtype-specific condition-2 ADA matrix. The number of reorganized TADs contributing to each subtype is indicated by *n*. The *strength-change up* and *strength-change down* groups show increased and decreased within-domain contacts, respectively, while boundary positions are retained. The *split* group shows broadly weaker within-domain contacts, particularly near both domain boundaries. The *zoom* group shows redistributed contacts within reference TADs and adjacent regions, whereas the *merge* group shows increased contacts with flanking regions. The *loss* group shows reduced contrast between within-domain and flanking contacts, consistent with weakened domain insulation, and the *complex* group shows spatially distributed contact changes. **b**, Corresponding ADA analysis of all reorganized TADs detected by DiffDomain-Spectrum, including those also classified as reorganized based on scHiCluster differential-boundary calls. Comparable subtype-consistent aggregate contact patterns were observed across the four representative comparisons. ADA was performed using `coolpuppy` on 25 kb aggregated scHi-C contact maps constructed from the developing mouse embryo HiRES dataset of Liu et al. [20].

**Supplementary Figure 3:**
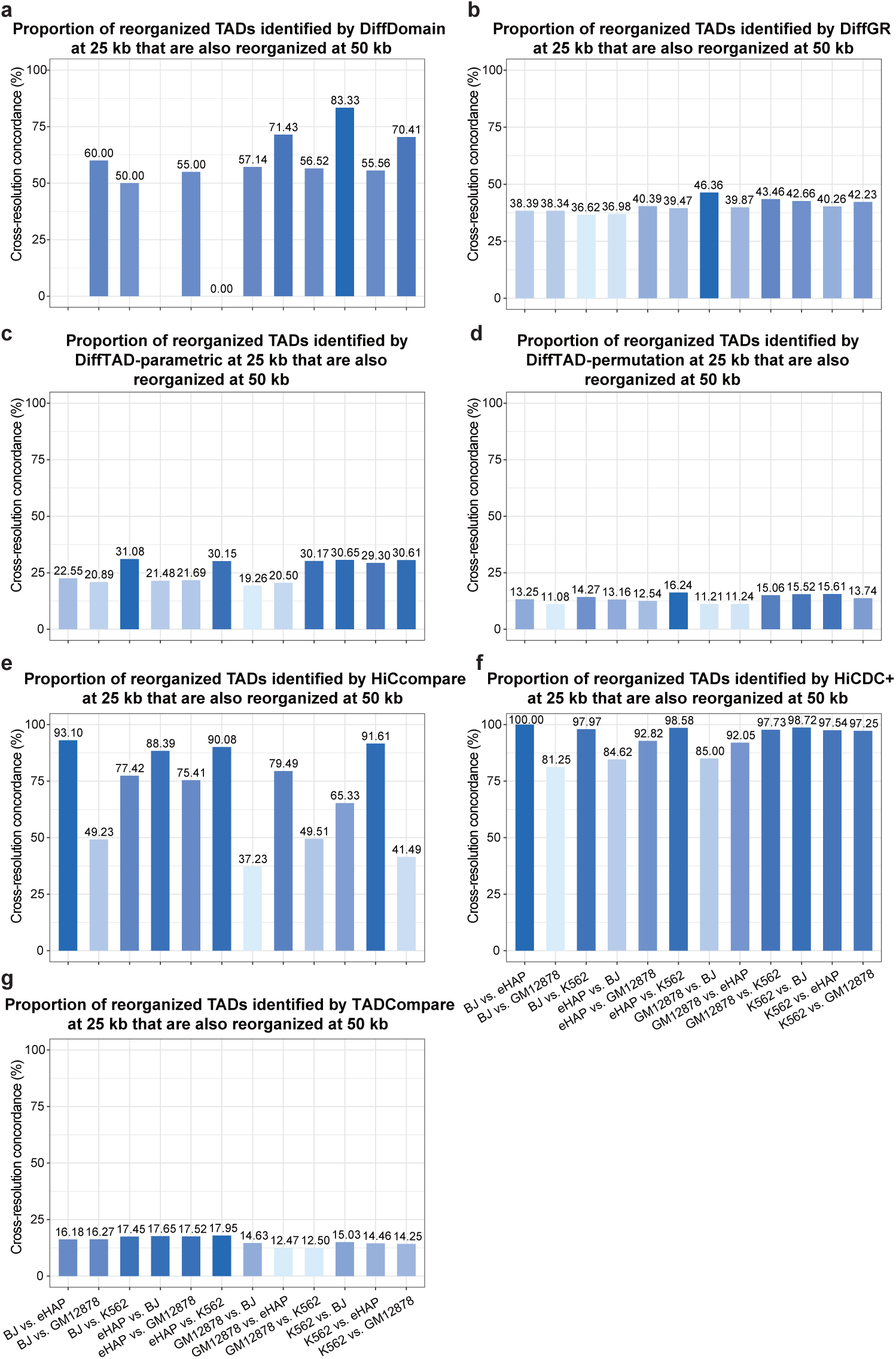
Cross-resolution concordance of reorganized TADs identified by seven alternative bulk callers. For each method, bars show the proportion of reference TADs detected as reorganized at 25 kb that were also detected as reorganized at 50 kb across 12 directed reference-TAD comparisons among GM12878, K562, BJ and eHAP. Panels show results for **a**, DiffDomain; **b**, DiffGR; **c**, DiffTAD-parametric; **d**, DiffTAD-permutation; **e**, HiCcompare; **f**, HiCDC+; and **g**, TADCompare. Concordance was at least 50% in nine comparisons for DiffDomain, eight for HiCcompare and all 12 for HiCDC+, but in none for DiffGR, DiffTAD-parametric, DiffTAD-permutation or TADCompare.

**Supplementary Figure 4:**
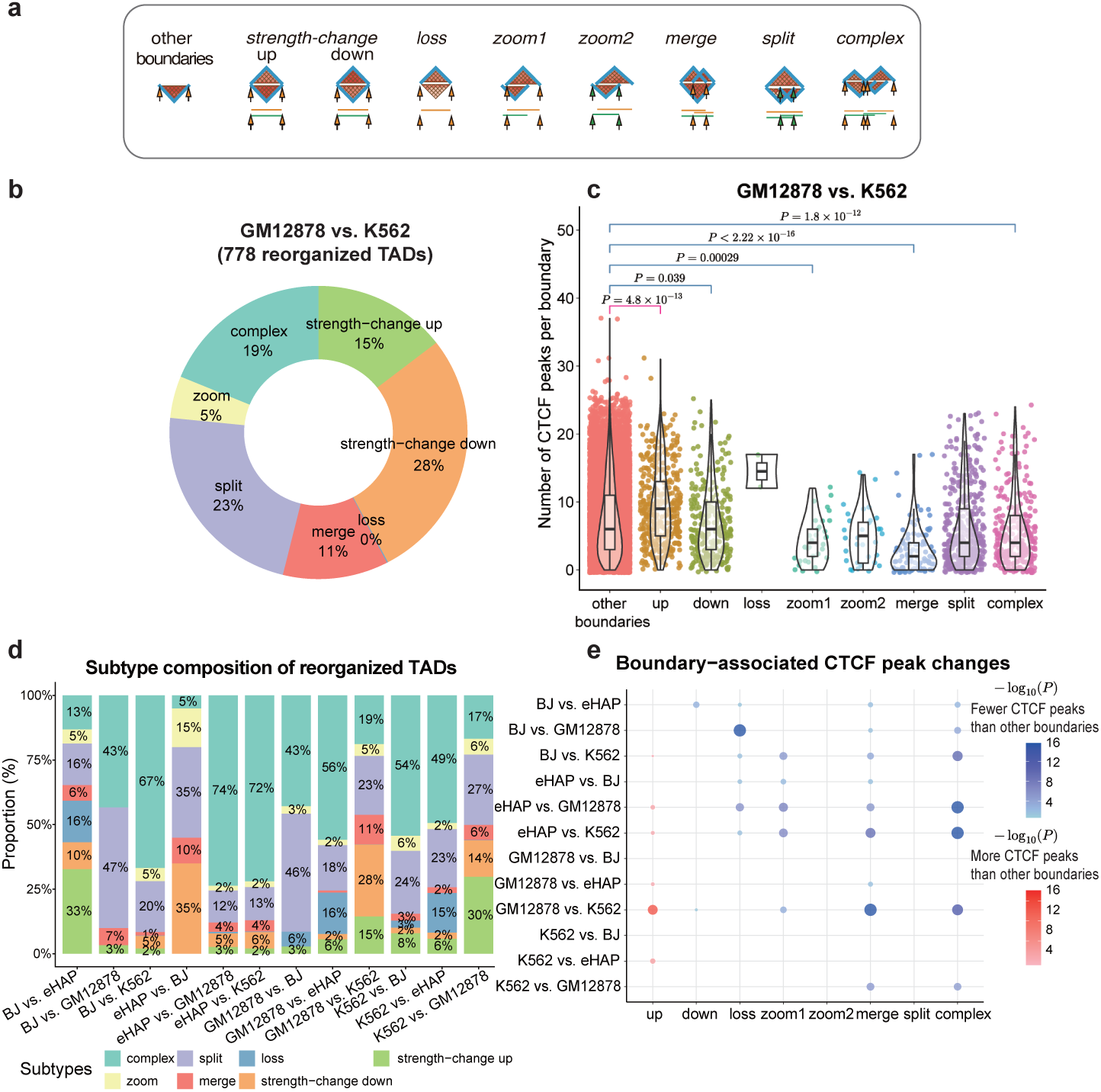
Subtype composition and boundary-associated CTCF peak changes in reorganized TADs detected by DiffDomain-Spectrum. **a**, Schematic definition of the boundary groups used for CTCF analysis. Condition-1 reference TADs and condition-2 TADs are shown above and below, respectively. Boundary groups were defined according to the structural subtypes assigned by DiffDomain-Spectrum: *strength-change up*, *strength-change down*, *loss*, *zoom1*, *zoom2*, *merge*, *split* and *complex*. The *zoom1* and *zoom2* groups represent boundaries lost from condition 1 and gained in condition 2, respectively; *merge* and *split* represent boundaries lost through TAD merging and gained through TAD splitting, respectively. The “other boundaries” control group comprises condition-2 TAD boundaries that do not overlap condition-1 TADs detected as reorganized in condition 2. Arrows indicate the genomic positions assigned to each boundary group. CTCF peaks were counted using condition-2 data for all groups. **b**, Subtype composition of the 778 GM12878 reference TADs detected as reorganized in K562 by DiffDomain-Spectrum. **c**, Distribution of CTCF peak counts across boundary groups for the GM12878-versus-K562 comparison, with GM12878 as condition 1 and K562 as condition 2. Each point represents one TAD boundary. CTCF peak counts in each subtype-associated boundary group were compared with those in the “other boundaries” control group using one-sided Mann–Whitney U tests. Only comparisons with *P <* 0.05 are annotated. Boxplot center lines denote medians, boxes denote interquartile ranges, and whiskers extend to the most extreme values within 1.5× the interquartile range. **d**, Subtype composition of reorganized TADs across 12 reference-TAD comparisons among GM12878, K562, BJ and eHAP. The four cell lines form six pairwise contrasts, each evaluated in both reference directions; the first-named cell line in each comparison provides the reference TAD set. **e**, Bubble plot summarizing differences in CTCF peak counts between subtype-associated boundary groups and the “other boundaries” control group across the 12 reference-TAD comparisons. Rows denote reference-TAD comparisons and columns denote the displayed subtype-associated boundary groups. Red and blue indicate significantly higher and lower CTCF peak counts, respectively, in subtype-associated boundaries relative to other boundaries. Bubble size represents − log_10_(*P*) from one-sided Mann–Whitney U tests. Comparisons with *P ≥* 0.05 are left blank.

**Supplementary Figure 5:**
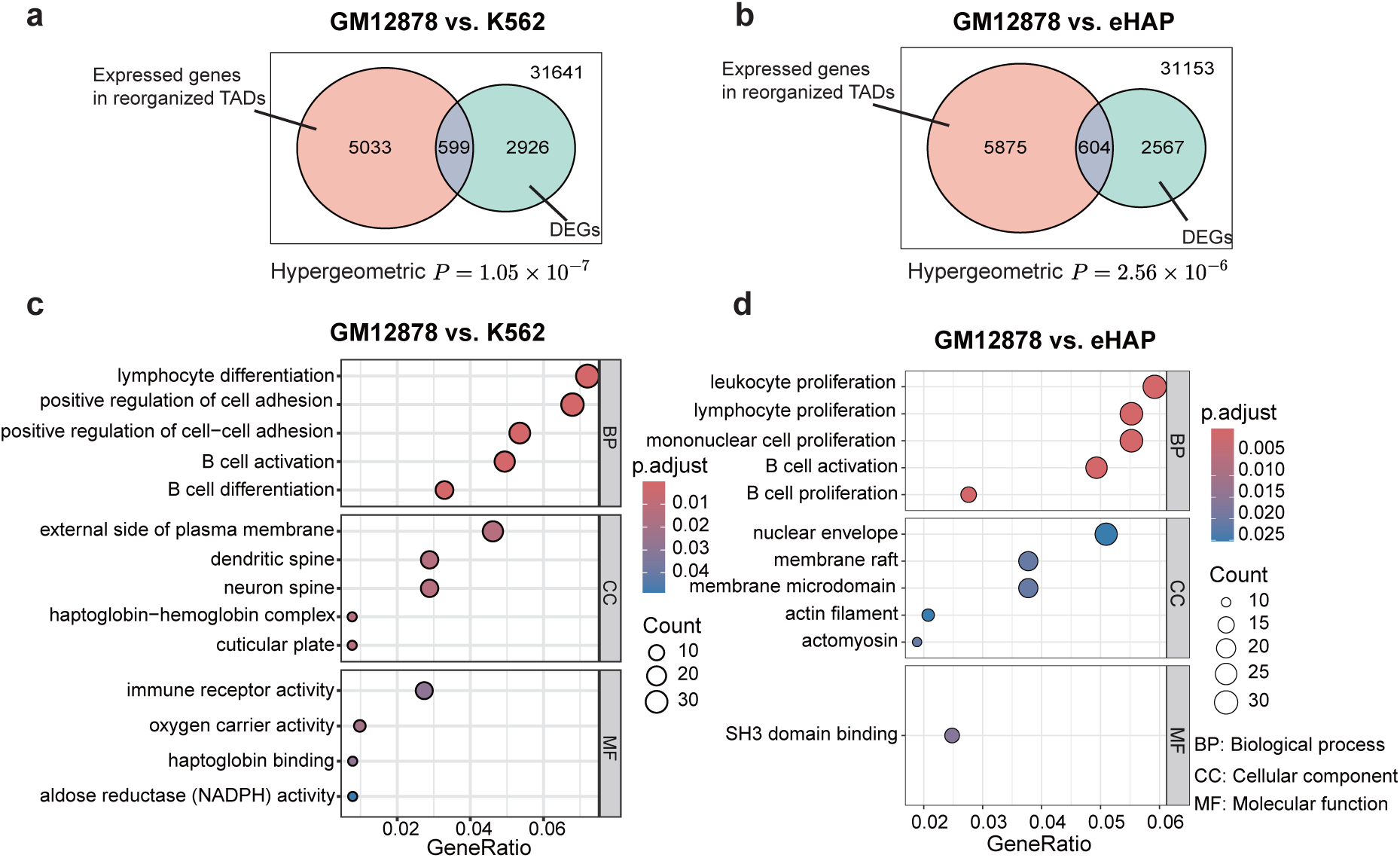
Differentially expressed genes are enriched within reorganized TADs detected by DiffDomain-Spectrum. **a,b**, Overlap between DEGs detected by DESeq2 from matched scRNA-seq data and expressed genes located within reorganized TADs detected by DiffDomain-Spectrum for **a**, GM12878 versus K562 and **b**, GM12878 versus eHAP. scRNA-seq and scHi-C were co-assayed in the same individual cells. The aggregated scHi-C contact maps comprised 221 GM12878 cells and 63 K562 cells in **a**, and 221 GM12878 cells and 42 eHAP cells in **b**. The overlaps contained 599 and 604 DEGs, respectively. Enrichment of DEGs among expressed genes located within reorganized TADs was assessed using hypergeometric tests, with genes overlapping any tested condition-1 reference TAD as the background set (*P* = 1.05 × 10*^−^*^7^ for GM12878 versus K562 and *P* = 2.56 × 10*^−^*^6^ for GM12878 versus eHAP). **c**, Gene Ontology enrichment analysis of the 599 genes overlapping between DEGs and expressed genes located within reorganized TADs in the GM12878-versus-K562 comparison. Selected significantly enriched terms are shown for biological process (BP), cellular component (CC) and molecular function (MF). Dot position indicates gene ratio, dot size indicates the number of overlapping genes assigned to each term, and color indicates the Benjamini–Hochberg-adjusted *P* value. **d**, Gene Ontology enrichment analysis of the 604 overlapping genes in the GM12878-versus-eHAP comparison. Selected significantly enriched terms are grouped by BP, CC and MF. Dot position indicates gene ratio, dot size indicates the number of overlapping genes assigned to each term, and color indicates the Benjamini–Hochberg-adjusted *P* value.

**Supplementary Figure 6:**
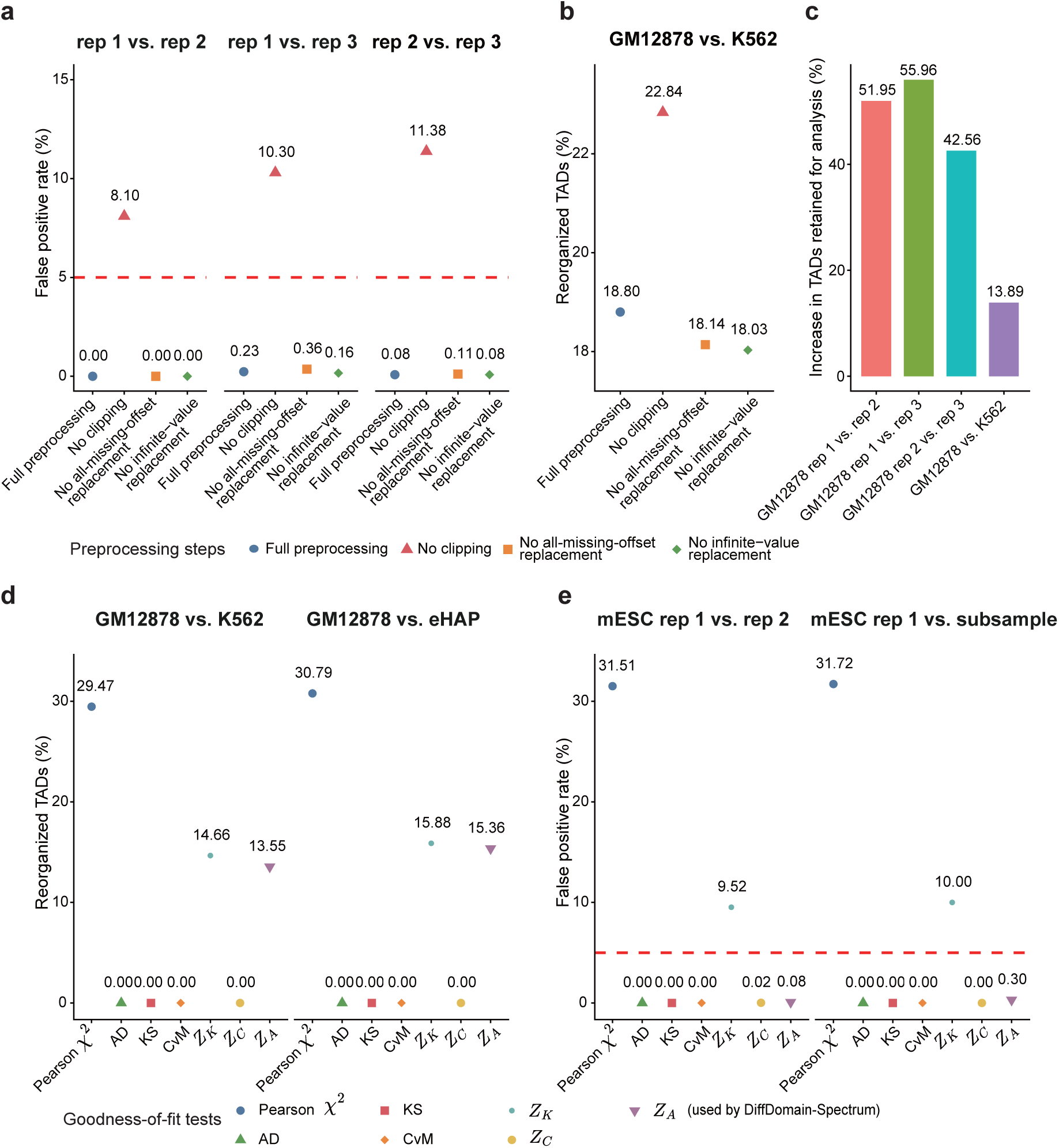
Ablation analysis of preprocessing steps and goodness-of-fit tests in DiffDomain-Spectrum. **a**, False positive rates (FPRs) across three pairwise comparisons among GM12878 scNanoHi-C2 biological replicates at 50 kb resolution. The evaluated preprocessing operations were applied to the log-scale TAD-level difference matrix rather than to the aggregated scHi-C contact maps. Full preprocessing was compared with variants omitting clipping of finite entries to [−3, 3], all-missing-offset replacement or distance-specific replacement of infinite entries. The dashed line indicates the nominal 5% FPR level. **b**, Proportion of reference TADs detected as reorganized in the GM12878-versus-K562 comparison using aggregated scHi-C contact maps generated by scNanoHi-C2 at 50 kb resolution under the same four preprocessing configurations. **c**, Relative increase in the proportion of TADs retained for analysis after all-missing-offset replacement compared with the corresponding analysis without all-missing-offset replacement, shown for three GM12878 biological-replicate comparisons and GM12878 versus K562 using scNanoHi-C2 data at 50 kb resolution. **d**, Proportion of reference TADs detected as reorganized by alternative goodness-of-fit tests in the LiMCA GM12878-versus-K562 and GM12878-versus-eHAP comparisons at 25 kb resolution. *Z_A_*denotes the full-spectrum goodness-of-fit test used by DiffDomain-Spectrum and is compared with Pearson’s chi-squared *χ*^2^, Anderson–Darling (AD), Kolmogorov–Smirnov (KS), Cramér–von Mises (CvM), *Z_K_* and *Z_C_* tests. **e**, FPRs of the same goodness-of-fit tests for a dscHi-C mESC biological-replicate comparison and a replicate-versus-subsample control at 25 kb resolution. The dashed line indicates the nominal 5% FPR level.

